# Unlocking the Full Potential of Nanopore Sequencing: Tips, Tricks, and Advanced Data Analysis Techniques

**DOI:** 10.1101/2023.12.06.570356

**Authors:** Daria Meyer, Winfried Goettsch, Jannes Spannenberg, Bettina Stieber, Sebastian Krautwurst, Martin Hölzer, Christian Brandt, Jörg Linde, Christian Höner zu Siederdissen, Akash Srivastava, Milena Zarkovic, Damian Wollny, Manja Marz

**Author notes:** These authors contributed equally (Daria Meyer, Winfried Goettsch, Jannes Spannenberg). These authors contributed equally (Damian Wollny, Manja Marz).

## Abstract

Nucleic acid sequencing is the process of identifying the sequence of DNA or RNA, with DNA used for genomes and RNA for transcriptomes. Deciphering this information has the potential to greatly advance our understanding of genomic features and cellular functions. In comparison to other available sequencing methods, nanopore sequencing stands out due to its unique advantages of processing long nucleic acid strands in real time, within a small portable device, enabling the rapid analysis of samples in diverse settings. Evolving over the past decade, nanopore sequencing remains in a state of ongoing development and refinement, resulting in persistent challenges in protocols and technology. This article employs an interdisciplinary approach, evaluating experimental and computational methods to address critical gaps in our understanding in order to maximize the information gain from this advancing technology. Here we present both overview and analysis of all aspects of nanopore sequencing by providing statistically supported insights. Thus, we aim to provide fresh perspectives on nanopore sequencing and give comprehensive guidelines for the diverse challenges that frequently impede optimal experimental outcomes.

## INTRODUCTION

Nanopore sequencing is a transformative technology in genomics, offering the unique ability to sequence DNA or RNA molecules in their native form. Nanopore sequencing can generate long sequencing reads by measuring disturbances in the ion current as biological molecules such as DNA and RNA pass through a nanopore (1, 2). During or after sequencing, the raw DNA/RNA signal can be transformed into nucleotide sequences. This capability provides invaluable insights into genetic variation and molecular modifications. Since the debut of the MinION device by Oxford Nanopore Technologies (ONT) in 2014, nanopore sequencing has experienced a surge in popularity and has become a cornerstone in genomic research. For DNA sequencing (2), it is attractive in diagnostics (3) and for metagenomic assemblies (4, 5, 6, 7), enabling the study of microbial communities and their functions. In direct RNA sequencing, it supports diverse tasks such as *de novo* transcriptome assembly, isoform expression quantification (8), and the direct detection of RNA modifications (9). These applications highlight the technology’s versatility and its ability to address a wide range of biological questions.

One of the key advantages of nanopore sequencing is its ability to generate data rapidly, facilitated by quick library preparation and real-time data acquisition during sequencing (10). Additionally, nanopore sequencing can directly sequence native RNA molecules without the need for reverse transcription or amplification (8). Its capacity for long-read sequencing, where fragments up to two megabases in length can be read in a single pass (11), further enhances its utility by providing comprehensive genomic and transcriptomic insights. Furthermore, due to their portability and affordability, devices such as the MinION (12) are particularly valuable for applications such as monitoring virus outbreaks, where fast and on-site sequencing is essential.

Nanopore sequencing offers many advantages but comes with technical challenges. When first introduced, the technology was limited by lower accuracy (∼60 %) and throughput, though these shortcoming have improved significatly over time (13). With the introduction of ONT’s MinION device in 2014 accuracy improved to 92 % (14), with current rates reaching 99 % (15). While in 2021 nanopore sequencing continued to exhibit a higher error rate compared to Illumina sequencing (2), in 2024 sequencing of nearly complete bacterial genomes without short-read or reference polishing became possible (16, 17). Nonetheless, sequencing accuracy remains a key consideration for certain high-precision applications, particularly in direct RNA sequencing (18).

Another challenge lies in the customization of nanopore wet lab workflows. While numerous reviews provide detailed overviews of tools for standard nanopore sequencing applications (10, 19, 20, 21), the field’s relative novelty has given rise to a broad range of non-standard approaches aimed at maximizing sample utility. The versatility of nanopore sequencing lends itself to customization to meet specific requirements, including the use of various flow cells, library preparation kits, sequencing buffers, and an extensive set of computational analysis tools. The sheer number of options can be daunting for newcomers. Optimizing protocols, designing libraries, and selecting parameters require careful consideration. Compounding this issue is the prevalence of unverified claims in the field, as many methods lack experimental validation, making them challenging to integrate into standardized workflows. This is particularly problematic when information is available solely through the Nanopore Community^1^ and lacks proper citation.

This perspective work integrates insights gained from over 300 nanopore sequencing runs, which span a variety of species, sequencing devices, preparation methods, and sequencing protocols. These data-driven findings provide valuable guidance for addressing common challenges in nanopore sequencing. For instance, we show that flow cell performance is influenced more by the sample type than by factors such as read length, flow cell age, or the number of active pores at the start of a sequencing run. Techniques such as flow cell washing and adaptive sampling (22) have been demonstrated to enhance output and improve sequencing yield. Additionally, we offer practical advice for optimizing library preparation, particularly for obtaining long reads and adjusting loading amounts when working with small sample sizes. We also address the complexities of data analysis, providing guidelines for constructing customized bioinformatics pipelines that align with specific experimental goals. Effective methods for calling RNA and DNA modifications, as well as strategies for normalizing raw signal data, are also discussed. By bridging the gap between experimental data and best practices, this review equips researchers with actionable strategies for optimizing nanopore sequencing experiments.

### BACKGROUND ON NANOPORE HARDWARE

At the heart of nanopore sequencing is a protein channel (nanopore) through which DNA or RNA molecules can pass (Fig. 1). These nanopores are sensitive to electrical changes as molecules move through them (23). Flow cells are filled with an electrolyte solution that establishes in absence of DNA or RNA a relatively constant electrical current through the nanopore. The sensor continuously records this current as a raw electrical signal in picoamperes (pA) (24, 25, 26). During library preparation of DNA/RNA, a motor protein is attached to the nucleotide strand, guiding the DNA or RNA to the nanopore and then unwinding it while controlling the pace at which the strand passes through, ensuring accurate sequencing (2). As each DNA or RNA molecule passes through a nanopore, it disrupts the electrical current in a unique way based on its sequence at a time (9 nt in the nanopore serve for the signal in flowcell R10 and 5 nt for the discontinued flowcell R9) (14, 27). The recorded electrical signal of the sensor, often referred to as the “squiggle”, transmits the data for downstream processing (10).

**Figure 1.**
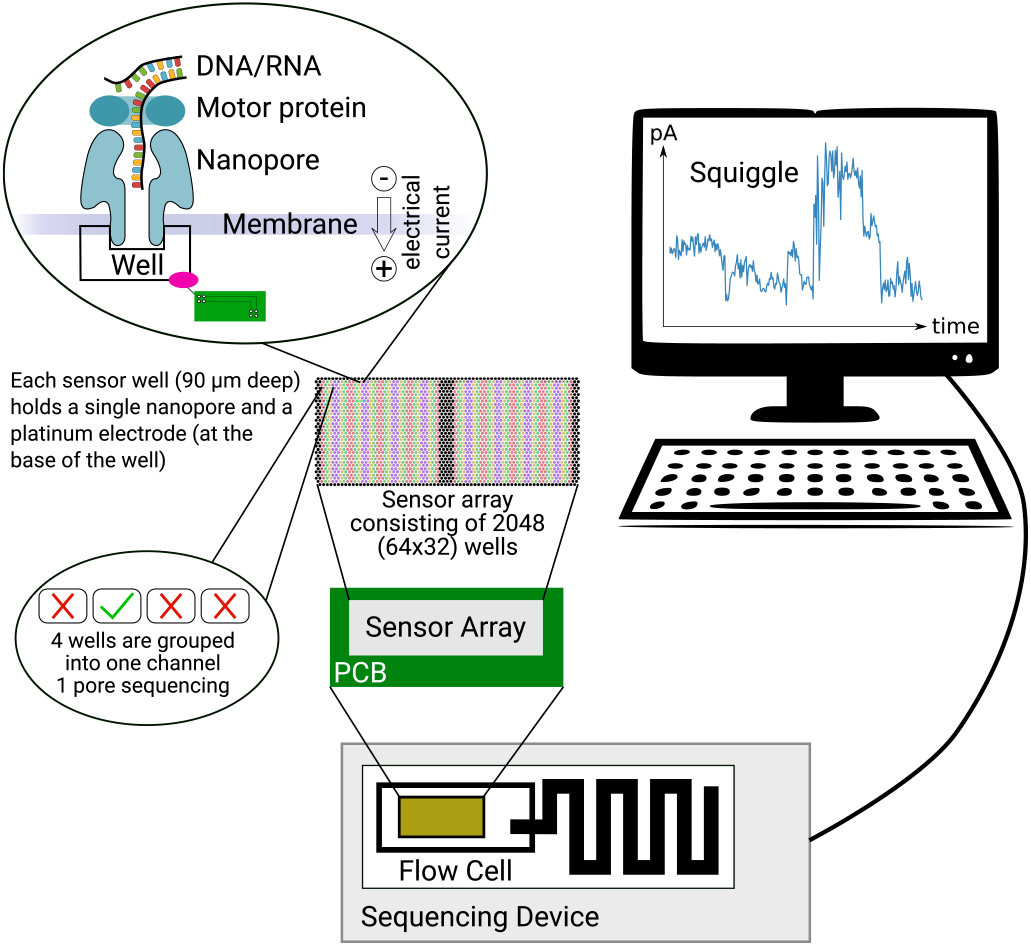
Nanopore sequencing on a MinION. A MinION flow cell contains 512 **channels** with 4 nanopores in each channel, resulting in 2 048 nanopores available for sequencing DNA or RNA. Each sensor well (90 *μ*m deep) holds a single nanopore and a platinum electrode (at the base of the well). The electrode is controlled via the **PCB** (Printed Circuit Board) and measures the ion flow as **ionic current**. During sequencing, a **motor protein** guides the DNA/RNA through the nanopore following the electric gradient. The DNA/RNA disrupts the current by partially blocking the ion flow. This can be measured in a resulting **squiggle**.

Each flow cell contains thousands of nanopores, which are grouped in channels of 4 pores. A standard MinION flow cell has 2 048 pores organized in 512 channels (2). Flow cells are the standard, full-sized sequencing units used in devices like MinION, GridION, and PromethION, which generate tens to hundreds of gigabases (Gb) of data (28).

Flow cells differ in the design of the nanopores, which impacts their performance mainly in terms of sequencing accuracy (29). In this work, we use R9 and R10 flow cells using single-constriction pores which were designed to be robust, reliable, and capable of handling a wide range of sequencing tasks.

The R10 flow cells feature double-constriction pores, where the nucleotide strand passes through two constriction points instead of one. The double-constriction design enhances the resolution of basecalling and can achieve higher raw sequencing accuracy (30).

A flow cell check can be performed manually in the `MinKNOW` software to assess the number of nanopores available in the flow cell before a sequencing run. At the start of each sequencing run an initial flow cell check is performed providing the starting number of available pores (31, 32). Each pore can only process a limited number of molecules before it becomes inactive. Therefore the number of available pores decreases over time (30, 31). Usually even a new flow cell never has the maximum number of pores available. A warranty is provided by ONT that allows flow cells to be returned if they do not have enough available pores within three months of purchase^2^.

The channels in the flow cell are multiplexed (mux), meaning that only a subset of nanopores can be actively used for sequencing at any given time. A mux scan identifies the most efficient nanopores on a flow cell by testing multiple groups of nanopores (mux groups) in different channels (33, 34). One pore per channel is selected every 1.5 h to continue sequencing within the channel, whereby the current pore can also be re-selected directly. The interval of 1.5 h is the default, but the time can be set in the `MinKNOW` software at the start of each sequencing run. This process helps to optimize the sequencing run by ensuring only high-quality pores are actively sequencing and increase the overall lifetime of the flow cell.

For more details we refer to the nanopore documentation^3^.

### DATA CHARACTERIZATION AND DESCRIPTION

This perspective draws its conclusions from an extensive and diverse dataset, encompassing over 300 nanopore sequencing runs Among these, a specific focus is placed on about 241 sequencing runs conducted using R9 and R10 MinION and GridION flow cells. This dataset offers robust insights into sequencing performance across various conditions. The study also acknowledges instances of sequencing failures and runs conducted on earlier flow cell types (e.g., flongles or R8 flow cells), although these are not included in the accompanying statistical analyses.

The 241 sequencing runs we focus on (see Tab. S1) include 207 DNA sequencing runs and 34 direct RNA sequencing runs. These runs span a wide range of organisms, including 33 virus, 45 bacteria, 5 protist, 2 plant, 47 insect, 17 mouse, 38 human, 51 metagenomic, and 3 synthetic samples. The amount of input material loaded onto the flow cells ranged from 42 ng to 1440 ng.

The output of sequencing runs was assessed based on several metrics: (1) **EB12** As the sequencing yield, defined as the total amount of sequenced bases, depends on the runtime of the flow cell, we decided to compare the total number of bases produced after 12 h We chose 12 h because some runs included in this study were stopped after that time to wash the flowcell. Referred to as EB12 in this study and estimated by `MinKNOW`, EB12 ranged from 7 · 10^6^ to 28 · 10^9^ nucleotides. For washed flow cells, only the data from the first run was included in the analysis of EB12. (2) The **median read length** calculated after basecalling varied significantly, ranging from 419 to 23 746 bases. We chose to calculate the median instead of the mean, as it is robust against outliers. (3) Another key metric was the **pore half time** of the flow cells, defined as the duration of sequencing until only half of the initially active pores remained active. This value ranged from 1 to 59 hours, highlighting variations in flow cell durability and performance across experiments. Due to artifacts observed in some flow cells, we manually re-evaluated the pore half time of 22 flow cells, as shown in Tab. S3. (4) Other flow cell parameters evaluated in this study include the number of active channels, active pores, and the age of the flow cells. The number of **active channels**, as reported in ONT run reports, ranged from 93 to 512 out of a possible 512 channels. (5) Similarly, the number of **active pores** varied between 749 and 1 984 out of a total of 2 048 available pores. As the number of active pores can increase shortly after starting sequencing, we used the maximum number of active pores during the first 10 minutes of each run. (6) The **flow cell age** was another important variable considered. This was calculated from the time of their arrival at the laboratory, as the production dates were not accessible. We analyzed flow cells that were used between 2 days and 249 days after their arrival, with ONT’s recommended maximum shelf life being 90 days when stored at 2–8°C. (7) Finally, we elaborated how the amount of loaded library (in ng) affects sequencing yield.

The following analyses provide deeper insights into flow cell performance and durability and have been conducted using custom scripts.

### CHALLENGES IN LIBRARY PREPARATION FOR ONT SEQUENCING

Library preparation for ONT sequencing involves multiple steps. For instance, DNA library preparation using the Ligation Sequencing DNA V14 kit (SQK-LSK114) entails DNA repair and end-preparation, followed by adapter ligation and clean-up, before priming and loading the prepared library onto the flow cell. An alternative, if short preparation times are a main focus, is the Rapid Sequencing kit V14 (SQK-RAD114) which reduces the preparation time to as little as 10 min at the cost of sequencing accuracy. For DNA, multiplexing is possible (35) using the Native Barcoding Kit 96 V14 (SQK-NBD114.96).

Similarly, library preparation for direct RNA sequencing (SQK-RNA004) begins with adapter ligation of the reverse transcription (RTA), followed by a RT reaction to stabilize the single RNA strand, and concludes with the ligation of the RNA Ligation Adapter (RLA), priming, and flow cell loading. Each stage of these protocols requires multiple handling and washing steps such as pipetting, mixing, and centrifugation, which introduce challenges in maintaining nucleic acid length and achieving consistent yields.

#### Obtaining long reads

To obtain the longest possible DNA or RNA reads, it is crucial to minimize any shearing of nucleic acids.

To avoid shearing during library preparation, it is important to reduce the suction force when pipetting. This can be minimized by using cut-tips and pipetting slowly, as recommended by Prall et al. (36). Additionally, vortexing is considered too harsh for nucleic acids, as it often results in shorter fragments, so it is preferable to use gentle tapping for mixing instead (37).

Many DNA/RNA isolation kits, particularly those using beat-beating- or column-based methods, are known to shear and fragment nucleic acids (38, 39). We recommend the classic phenol-chloroform extraction method, as it preserves longer fragments. Recently, there has been a growing interest in high molecular weight DNA extraction techniques. Innovations in this area include the surface topography of silica lamellae and new solid-phase methods introduced by different brands, which aim to improve yield and ease of handling. These kits are often more user-friendly but typically come at a higher reagent cost (40, 41).

In our analysis of DNA and RNA sequencing reads, we mainly used slow and gentle phenol-chloroform extraction to avoid nucleic acid shearing and to maximize read length. We achieved notably long reads in both DNA and RNA samples. The longest DNA read obtained was 1.4 Mb, which came from a metagenomic sample (sample ID: 117.1) and was classified as *Homo sapiens* using `kraken2` (42), with a GC content of 41.4 %. While nanopore sequencing is generally more robust in handling GC-rich regions than PCR-dependent platforms, high GC content can still affect read quality, base-calling accuracy, and sequencing coverage (43, 44). For RNA, the longest read was 175 kb from sample ID 103.1, which classified as bacterial with a GC content of 68.6 %.

#### Adjustment of loading amount

##### Loading dependent on pore availability

While it may seem intuitive to increase the amount of DNA/RNA loaded to enhance data yield, there is a threshold beyond which additional loading may not yield proportional benefits^4^. This is particularly true due to saturation effects caused by a limited number of pores on the flow cell. The quantity of data generated is directly proportional to the initial number of active pores (Fig. 2A) and channels (Fig. 2B). When working with flow cells that initially have fewer pores available, it is advisable to load less sample material. Our experiments demonstrated that for flow cells with at least 1 500 active pores, a DNA load of 350 ng produced reasonable sequencing output. For flow cells with approximately 1 200 active pores, we made experience with 300 ng of DNA, and for those with around 800 active pores, we loaded about 250 ng, as detailed in Tab. S1. However, the amount of loaded library does not correle (R^2^ - 0.08) with the resulting EB12 for DNA (see Fig. 2C). A relatively weak trend towards higher EB12 is observed when loading more starting material for RNA (R^2^ - 0.20).

**Figure 2.**
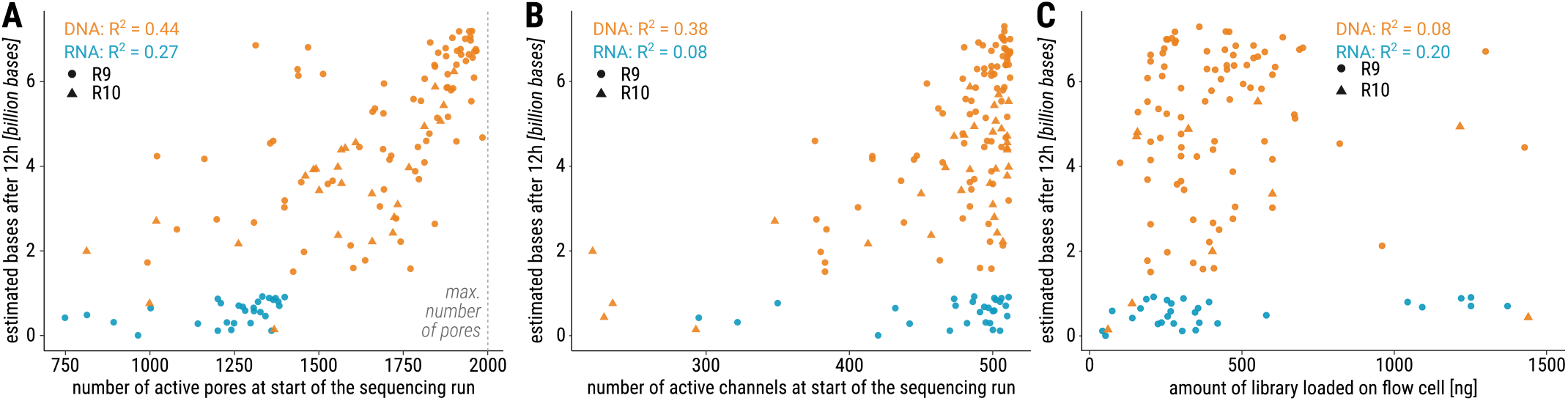
**(A)** The number of active pores (maximum number of active pores during the first 10 minutes) at the beginning of the sequencing run seem to influence EB12. Note the generally lower number of active pores for RNA runs. **(B)** The number of active channels at the beginning of the sequencing run has a weaker influence on EB12. Although a high yield is only achievable when many channels are active, starting with many active channels does not automatically guarantee a high EB12. R^2^ values were calculated for second degree polynomial regression. For DNA a correlation can be seen between estimated bases after 12 h and amount of pores and channels at the sequencing start, respectively. For RNA, a relatively weak trend can be seen only for the amount of active pores. **(C)** For DNA, the amount of estimated bases after 12 h is independent of the amount of starting material loaded onto the flow cell (R^2^ = 0.08). For RNA, a relatively weak trend shows increased amount of estimated bases after 12 h with increased amount of starting material loaded onto the flow cell (R^2^ = 0.20).

Note that R10 flow cells exhibit similar behavior in this regard as the discontinued R9 flow cells.

Instead, having extra input material available is advantageous for repeated washing of the flow cell, as it helps recover nanopores and extend overall sequencing yield. Specifically, when planning to wash a flow cell, we recommend splitting the library for different sequencing runs in advance, see details in Sec. Washing and reusing flow cells. **Minimal DNA loads for flow cells** For optimal results in DNA sequencing, ONT recommends to start the library preparation with 1000 ng (equivalent to 100–200 fmol) of total DNA (SQK-LSK114 DNA library preparation kit^5^). Following library preparation, approx. 50 % of the DNA is typically available for loading into the flow cell.

For samples with limited DNA quantities that cannot be amplified, the following methods have yielded successful results: Although ONT’s instruction recommend a minimum of 100 ng^6^, studies show that without special treatment, 1 ng of DNA (prior to library preparation) yielded 6,118 reads with an N50 of 3,907 bases (45). This result indicates good quality; however, reduced input correlates with decreased output. Carrier sequencing allows for the sequencing of very small amounts of target DNA by mixing it with a larger quantity of non-target “carrier” DNA, ensuring adequate representation without amplification. This technique successfully detected as little as 0.2 ng of target *Bacillus subtilis* DNA combined with 1 000 ng of carrier Lambda DNA, producing high-quality data without amplification (46).

The 10X Genomics Chromium Controller is an advanced microfluidics-based platform that enables high-throughput single-cell and spatial analysis of DNA, RNA, and proteins. DNA extracted from the Chromium Controller can be sequenced on nanopore sequencers enhancing the analysis of low-input samples (47). The technology enables sequencing of genomic DNA quantities as low as 50 pg.

In our hands, when loading as low as 30 ng DNA onto a flow cell (435 ng before library preparation, ID 67.6) we successfully got 648 Mb data. In another run we loaded 50 ng onto the flow cell (ID 129.2), which resulted in even 1.5 Gb data output. Simon *et al*. reported the successful use of 50 ng in metagenomic samples with eight species (48).

##### Minimal RNA load for flow cells

RNA sequencing requires poly-adenylated 3’ ends (poly-A tails) that serve as consistent binding sites for ligating adapter molecules to create a continuous RNA-adapter complex that can pass through the nanopore, see Sec. Adapter ligation. Ribosomal RNA (rRNA), which lacks poly-A tails, makes up the majority of cellular RNA; for instance, *E. coli* cells contain around 85 % rRNA (49). Ribosomal RNA is naturally excluded when using a poly-A-based sequencing approach. Usually, this is intended, given that rRNA sequencing is often undesired. For optimal RNA sequencing, ONT recommends to use 500 ng of total RNA or 50 ng of poly-A-tailed RNA (which is in agreement with the ratio of rRNAs in the cell).

Many important RNA molecules, including certain mRNAs (e.g., histone-coding) and most polymerase III transcripts (like ncRNAs, non-canonical miRNAs, and lncRNAs), lack poly-A tails (50, 51). To sequence all RNAs except rRNAs, it is recommended to ligate poly-A tails to the sample and perform an rRNA depletion step prior to poly-A ligation (52). A method for total RNA sequencing with only 10 ng input material has been reported (53). In our hands, 42 ng RNA (ID 152.1) produced 157 Mb of data.

#### Using external control sequences

The use of external control sequences is a critical aspect of controlling library preparation and benchmarking the sequencing process in nanopore workflows. Both, the DNA ligation sequencing kit (SQK-LSK114) and the direct RNA sequencing kit (SQK-RNA004) contain positive control strands (spike-ins). The DNA Control Strand (DCS, Fig. 3) is a 3.6 kb standard amplicon that maps to the 3’ end of the Lambda virus genome. It is provided at a concentration of 10 ng/*μ*L.^7^.

**Figure 3.**
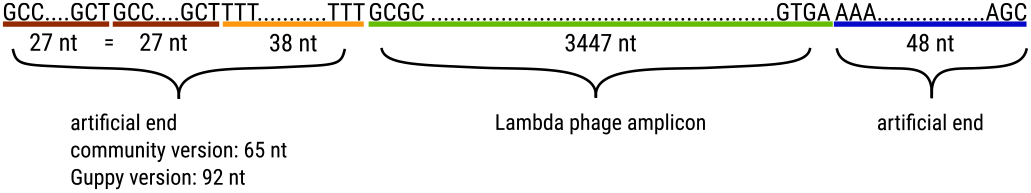
Structure of the ONT DNA spike-in: The positive control spike-in for DNA sequencing is a 3.6 kb amplicon, which maps to the 3’ end of the Lambda phage genome.

The RNA Calibrate Strand (RCS) is derived from the *Saccharomyces* genome, specifically the Enolase II gene (ENO2, YHR174W). It is supplied at a concentration of 15 ng/*μ*L^8^. Both DCS and RCS are utilized as positive controls for the library preparation and sequencing process; and therefore are not used for normalization of sequencing results. In our experiments, we detected approx. 1 % of DCS/RCS sequences in our samples.

#### Adapter ligation

Adapter ligation, the final library preparation step for nanopore sequencing, attaches adapters with motor proteins to sample molecules. These adapters guide DNA or RNA into the nanopore and regulate sequencing speed, making ligation efficiency crucial for optimal output. The concentration of the adapters in the mixes is proprietary; however, it is standardized for reactions involving approx. 100-200 fmol (approx. 1 *μ*g of DNA^9^), 50 ng for poly(A)-tailed RNA, and 500 ng for total RNA. Although the impact of RNA modifications on adapter ligation efficiency is not yet well investigated, these modifications remain an essential factor to consider for RNA samples (8).

New England Biolabs (NEB) generally recommends maintaining a 10:1 adapter-to-sample ratio to ensure efficient ligation without excessive free adapters^10^. According to ONT, high adapter-to-sample proportions generally do not hinder sequencing as long as the majority of pores are actively sequencing^11^

#### Choosing between LFB/SFB buffer

The buffer choice in the final library prep step determines DNA fragment retention. Long Fragment Buffer (LFB) enriches fragments ≥3 kb for long-read sequencing, while Short Fragment Buffer (SFB) retains a broad size range, affecting sequencing yield and flow cell performance. Our observations indicated that using SFB during the library preparation resulted in more sequencing output (EB12) when compared to the use of LFB (Fig. 4), which also applies to the estimated number of sequenced bases after 24 h and 36 h (see Fig. S3). These differences are evident when analyzing both flow cell types, R9 and R10, together. However, when analyzing R10 flow cells only, the difference between the buffer types was not as pronounced, Fig. S8. In the case of R9 data only, a slightly larger difference emerged between the outcomes of LFB and SFB applications, Fig. S9. These findings suggest that buffer choice may have a variable impact depending on the flow cell generation, potentially due to differences in pore design and chemistry.

**Figure 4.**
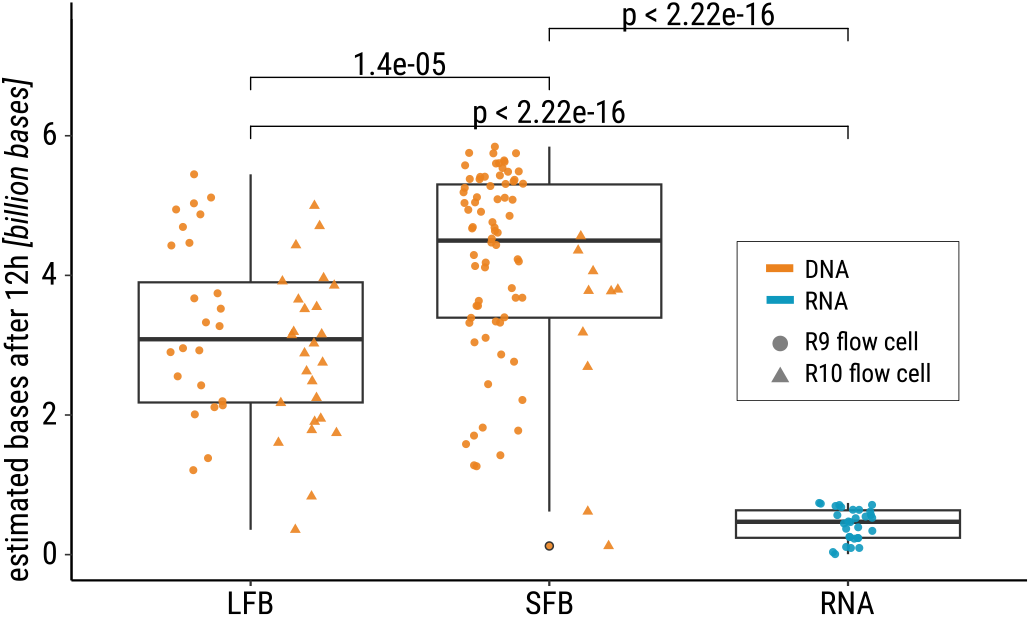
When combining R9 and R10 data, there are significant differences between the used buffers. As expected, RNA runs show significantly different amount of estimated bases after 12 h, but also the usage of LFB or SFB results in significantly different amount of sequenced bases after 12 h. For separate plots see Supplement Fig. S9 and Fig. S8. The finding remains unchanged for estimated bases after 24 h and 36 h, see Fig. S3.

#### Amplicon sequencing

Amplicon sequencing is one common strategy to enrich low-concentrated nucleic acid target material from complex samples for subsequent sequencing. Amplicon sequencing enriches specific DNA regions by amplifying them via PCR with specific primers. The resulting PCR products, or “amplicons,” are then sequenced to provide high-resolution information on the targeted regions. A key example of amplicon sequencing is its use in viruses, such as SARS-CoV-2 during COVID-19, for cost-effective sequencing and genomic surveillance. (54). ONT has specific advantages in sequencing long amplicons without additional fragmentation, *e*.*g*. to retain information of co-occurring mutations on single amplicons supporting virus variant deconvolution and lineage assignment (55).

In nanopore sequencing, amplicons should be mixed in equal nanomolar concentrations (56, 57, 58). When working with amplicons of the same size, mass concentration (nanograms per microliter) can be used instead of molar concentration. Due to the specific target enrichment, amplicon sequencing can ensure sufficient sequencing depth to reliably detect genetic mutations, supporting detailed genomic analysis in fields requiring accurate variant detection.

Amplicon sequencing can be an effective approach to explore entire virus genomes with low viral titers. While a lot of work was done on SARS-CoV-2, amplicon designs are increasingly being explored for other viruses, such as PPV. High-quality assemblies require overlapping amplicons, as accurate strain identification is crucial for diverse pathogens (see Fig. 5A). In some protocols, however, such as the widely used `ARTIC` protocol^12^ for SARS-CoV-2 sequencing, smaller amplicons of 392 bp are employed, with an overlap of 90 bp. In our study of SARS-CoV-2, we achieved high-quality results using longer amplicons of approx. 1200 bp, with an overlap of 117 bp between consecutive fragments (59). We demonstrate amplicon sequencing for Plum pox virus (PPV), which exhibits substantial genetic diversity across strains (Fig. 5A). We gained a uniform distribution of coverage across the entire genome (see Fig. 5B), demonstrating the effectiveness of well-designed amplicons for comprehensive genomic representation. Monitoring amplicon coverage is essential, as the aforementioned diversity may result in poor amplicon performance due to insufficient primer binding in mutated regions.

**Figure 5.**
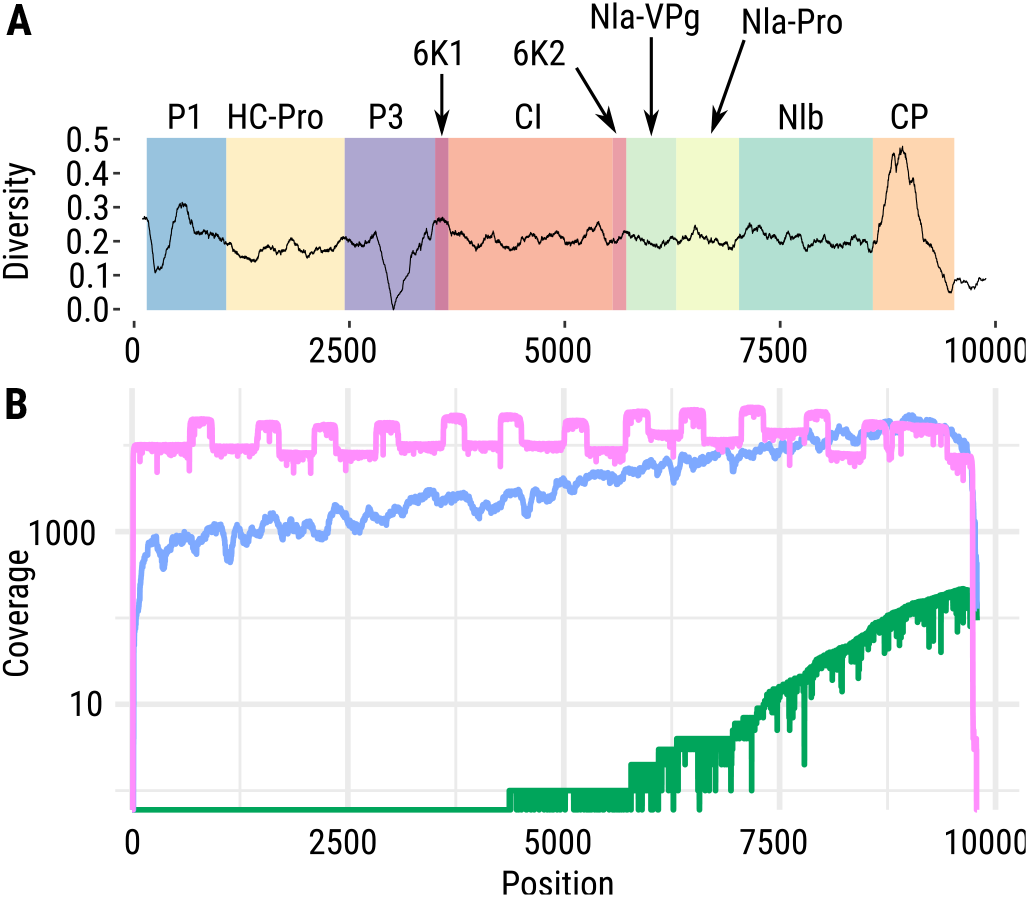
**A** Genetic diversity for each position in the plum pox virus (PPV) genome across 109 PPV isolates of all known PPV strains reduced in the P3 gene region and enhanced in the genetic region encoding for the capsid protein (CP). **B** Coverage comparison of three sequencing methods: direct RNA sequencing using nanopore (green), Illumina shot-gun RNA-Seq (blue) and nanopore-based PCR amplicon cDNA sequencing (pink) at the example of PPV. A characteristic feature of amplicon sequencing is the clear visibility of higher coverage regions due to the overlapping amplicon design and specific enrichment

#### Direct RNA Sequencing

The direct RNA sequencing protocol provides an efficient approach for sequencing RNA molecules directly without the need for converting them into complementary DNA (cDNA). This method allows for sequencing RNA in its native form, preserving RNA modifications, without amplification, and potentially capturing full-length RNA transcripts. An optional library preparation step, binding the RNA strand to a synthesized, complementary DNA strand, can increase throughput and improve read quality by unfolding complex RNA structures, which aids in producing more consistent sequencing data. However, this approach has downsides: (i) Prolonged incubation of RNA at high temperatures with divalent cations during cDNA synthesis can cause RNA degradation; (ii) Common reverse transcriptases may cleave RNA strands. This can reduce the availability of full-length RNA and impacting the efficiency and accuracy of direct RNA sequencing.

A distinctive feature of ONT’s direct RNA sequencing is its ability to detect RNA modifications. However, the bioinformatics analysis of these modifications is still underdeveloped. In this work, we focus intensively on this aspect, see Sec. Hidden treassures in raw data.

### TECHNICAL CHALLENGES IN NANOPORE SEQUENCING

#### Maximizing sequencing yield

Sequencing yield, also referred to as throughput, represents the total amount of data generated during a sequencing run. According to Wang *et al*. (2), the expected sequencing yield of a flow cell primarily depends on three factors: the number of active nanopores, the DNA/RNA translocation speed through the nanopore, and the running time. If the translocation speed is too fast, the system may struggle to differentiate between nucleotides, resulting in inaccurate sequencing. Conversely, if the translocation speed is too slow, throughput is reduced. The translocation speed is largely dependent on the motor protein used. For DNA, it typically operates at around 400-450 nt per second, while for RNA, the speed is slower, at approx. 70 nt per second (2). This speed can be monitored in real-time if live basecalling is enabled, although it can only be controlled indirectly. For example, increasing the temperature may accelerate the translocation speed, whereas overloading the flow cell might lead to a reduction in speed.

To analyze sequencing yield we explore over 300 ONT sequencing experiments across a variety of species, see Sec. Data characterization and description, Tab. S1. We observed significant variation in sequencing yield and investigate potential reasons for this yield disparity (Fig. 2, Fig. 6). We first investigate whether EB12 is influenced by the pore half time, which refers to the duration after which half of the nanopores in a sequencing run become inactive. A strong correlation is observed between pore half time and EB12 across all sample types except bacteria (Fig. 6A), consistent for both R9 and R10 flow cells This correlation remains significant even when normalizing EB12 for the number of starting pores, as shown in Fig. S4. As expected, a slower degradation rate of the flow cell leads to a higher sequencing yield.

**Figure 6.**
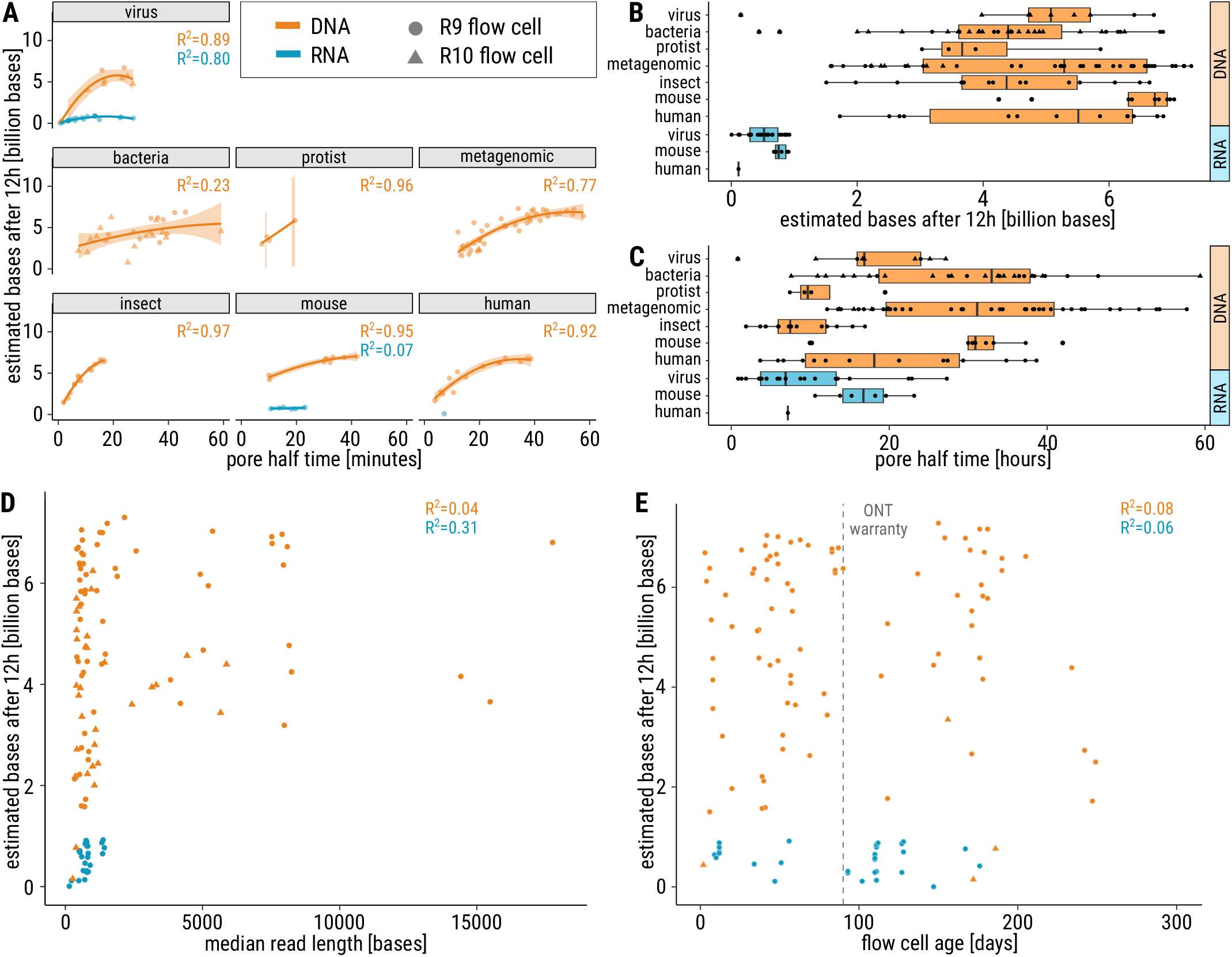
Maximizing sequencing yield. Sequencing yield from R9 (dots) and R10 (triangle) for DNA (orange) and RNA (blue) samples from a variety of species. Differences between RNA and DNA are evident in all plots, with a lower sequencing yield for RNA in general. Note, that the lower RNA yield is dependent on the slower sequencing speed (2) and the generally low number of active starting pores during RNA runs. **(A)** Correlation (*R*^2^) between pore half time and the estimated number of bases after 12 hours of sequencing (EB12) is observed across most sample types. No significant correlation is evident for bacteria due to high variability. Regarding R9 and R10 data separately, the correlation for bacteria slightly increases (R9: R^2^ = 0.39, R10: R^2^ = 0.11), but still remains weaker than for the other sample types. For the other sample types the correlation does not change when separating R9 and R10 flow cell data. **(B)** The EB12 yield is consistent across all sample types and flow cell type. **(C)** In contrast to EB12, the distribution of pore half time differs strongly between the sample types. Insect samples have a lower pore half time than other samples. Metagenomic and bacteria samples show a high variance. Data shown in B and C indicate differences between R9 and R10 data for metagenomic samples, see Fig. S10. **(D)** The median read length does not influence EB12 (nor the amount of estimated bases after 24 h and 36 h, see Fig. S7) for DNA. For RNA, a weak positive correlation is visible, which is slightly stronger after 24 h and 36 h, see Fig. S7. But note that there are only a few data points for RNA after 24 h and 36 h, and the amount of starting pores for RNA was generally lower. **(E)** The age of the flow cell (number of days from arrival to usage) does not affect EB12, although our data for R10 flow cells are limited.

Next, we analyze the impact of sample type on EB12. DNA sequencing yields in our data are generally higher than the ones for RNA, as shown in Fig. 6B, likely due to increased translocation speed and higher number of starting pores. Although DNA mouse samples have a slightly higher EB12 than other sample types with p-values of 0.04 or below (except p = 0.067 for viruses), see Fig. S5, it appears that after 12 hours of sequencing, the sequencing yield is roughly similar, regardless of the sample type.

Contrary, the pore half time seems to vary significantly depending on the type of sample being sequenced (Fig. 6C, with a median pairwise p-value of 0.011, see Fig. S6). For DNA samples, for instance, we found that insect and protist samples tend to have a much shorter pore half time than mouse, metagenomic, or bacterial samples. Furthermore, flow cells used for bacterial, metagenomic, and human samples exhibited high variability in pore half time. The flow cell pore half time appears to be influenced not only by RNA/DNA characteristics but also by the sample type. We assume these differences might originate from factors such as the quality and purity of RNA/DNA, the extraction method, the sequencing kit used or the cell type composition. Further research is needed to investigate those factors.

Despite variations in pore half time, the distribution of EB12 remains relatively consistent across different sample types (Fig. 6B). Notably, insects and protists, which exhibit shorter pore half time, do not stand out as outliers regarding EB12. When comparing R9 to R10, as shown in Fig. S10, there are no significant differences across all sample types. However, a significant reduction (*p<* 2.4 · 10^−7^)in both EB12 and pore half time is observed for R10 when analyzing metagenomic samples specifically. Taken together, after 12 hours, the sequencing output remains relatively consistent regardless of sample type, pore count and flow cell type, but the sample type seems to influences the degradation rate of the flow cell.

The median read length had no impact on EB12, see Fig. 6D, nor the amount of estimated bases after 24 h and 36 h, see Fig. S7. Most runs had a median read length below 3 kb, with EB12 values ranging from 0 to nearly 8 Gb and showing a fairly even distribution. Runs with a median read length exceeding 3 kb tended to have higher EB12 values, highlighting a clear distinction between RNA and DNA. R10 median read lengths aligned with those of R9 data.

We then turned our attention to the properties of the flow cells themselves. The age of the flow cell (the time between delivery and use) did not significantly affect EB12. Surprisingly, flow cells over 200 days old (Fig. 6E) performed similar to younger ones, although exceeding the ONT warranty period of three months by a factor of two. The oldest flow cell we tested was 249 days old, performing normally (1080 pores, 3.1 Gb output), see Tab. S1, ID 172.1.

In summary, a high EB12 is only achievable when many channels/pores are active at the beginning of the sequencing run (Fig. 2A,B). However, starting with many active channels does not automatically guarantee a high EB12. On the other hand, as expected, the number of active pores at the beginning of a sequencing run influence EB12, particularly when sequencing DNA (Fig. 2A).

#### Washing and reusing flow cells

Flow cells naturally degrade with use as active pores decrease, often due to blocking during sequencing. Blocked pores shift from “single pore” to “unavailable” in `MinKNOW`. While ONT acknowledges pore blocking, reasons for its variability remain unclear. Pores can be unblocked using voltage reversal during mux scans or by washing the flow cell.

Washing is performed with ONT’s “Flow Cell Wash Kit” (EXP-WSH003, EXP-WSH004), which uses DNase I to digest and remove loaded genetic material, freeing blocked pores. The cost of washing is minimal, approx. C16 per wash step, as the Wash Kit EXP-WSH004 costs C95 for six wash steps^13^. The sequencing run can be paused or stopped before washing: pausing allows continuation in the same file, while stopping generates a new file for subsequent sequencing. After washing, the flow cell can be reused for sequencing a new sample, but the previous run must be stopped rather than paused. Washed flow cells can also be stored for future use. However, they will not perform as well as new ones; the number of active pores will be lower than initially but higher than before washing (see Fig. 7).

**Figure 7.**
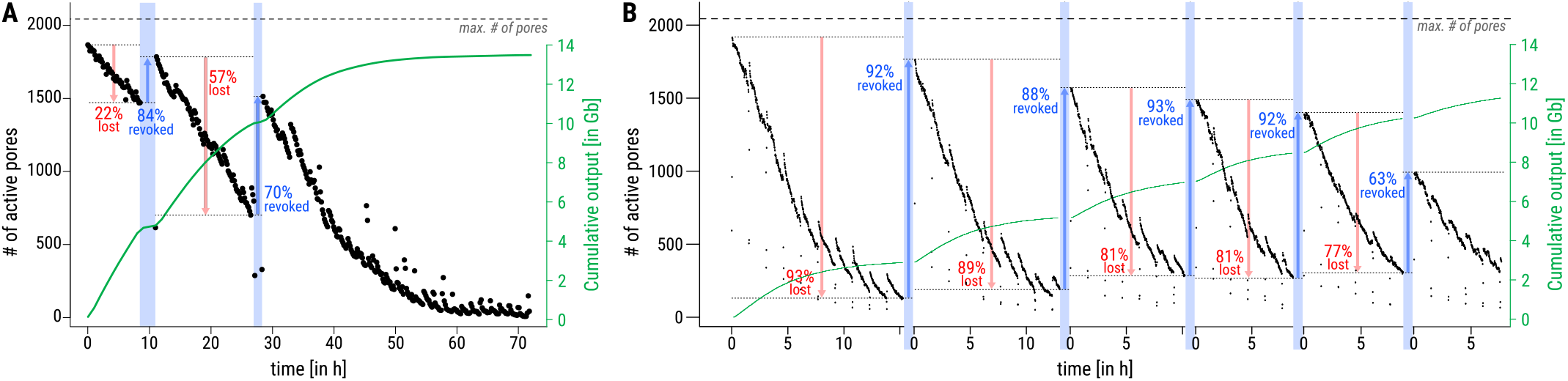
Influence of washing the flow cell on the amount of active pores. **(A)** A flowcell which has been paused (blue bars) two times for washing. **(B)** A flowcell which has been stopped (blue bars) five times for washing.

Washing can also be employed during a single sequencing run to increase sequencing yield by reactivating blocked pores. We identified three key factors to consider before sequencing: (i) potential washing steps, (ii) optimal washing timing, and (iii) available sample material. We investigated the impact of washing flow cells on pore recovery by analyzing 34 flow cells that had undergone 1-6 washes, totaling 58 washing steps, see Tab. S1. Examplarily, Fig. 7 shows two such washing steps. Statistics of these two flow cells are displayed in Tab. S4. In Fig. 7A, human DNA sample (ID 111.1) was washed twice during sequencing (at 10 h and 30 h), pausing the run (blue bars). Active pores decreased over time (red arrows) but increased after each wash (blue arrows). In Fig. 7B, the cricket DNA sample (ID 124.1–124.6) was washed five times during sequencing. A mean loss of 84.2 % of active pores could be compensated by revoking on average 85.6 % of those pores. ONT recommends washing and reusing a flow cell up to four times. Our findings show that flow cells can be washed at least five times and still produces data.

A key factor for high sequencing yield is determining the appropriate time point for washing, as yield depends on the number of active pores, Fig. 2A. It seems that pores do not block faster after washing. Therefore, we recommend monitoring cumulative output in `MinKNOW` : when the curve flattens (Fig. 7A, green line), we advise washing.

To ensure sufficient material for reloading, we recommend splitting the sample before starting the sequencing run if washing is planned to increase yield. For instance, when planning two wash steps, we recommend allocating 50 %, 30 %, and 20 % of the total library for the initial run and subsequent washes, respectively. This decreasing allocation accounts for the reduction in active pores after each wash. Adding excess material beyond the capacity of available pores will not further increase sequencing yield (see Fig. 2C and Fig. 7)

#### Adaptive sampling

Adaptive sampling allows for real-time selection or rejection of DNA fragments based on their sequence content, enabling targeted enrichment or depletion of specific genomic regions, organisms, or sequence types during a sequencing run^14^ (60). For example, adaptive sampling can enrich gene panels or exomes (61), eliminate unwanted sequences such as highly repetitive regions or contaminants, and target specific organisms or deplete host in metagenomic studies (22, 62, 63). This approach enhances coverage of targeted genes or regions without requiring complex library preparation steps like amplification or hybridization probes.

As DNA molecules pass through a nanopore, the device generates real-time sequence data (Fig. 8A). This requires a GPU and a fast basecalling model for adaptive sampling. The system quickly compares each fragment’s partial sequence (approx. 400 bases, corresponds to one second of data) to a reference genome or user-defined targets. When enriching for regions of interest, the sequencing continues only if the sequence matches the region of interest; otherwise, the DNA fragment is ejected, conserving flow cell resources and improving yield for desired regions. The depletion mode works vice versa.

**Figure 8.**
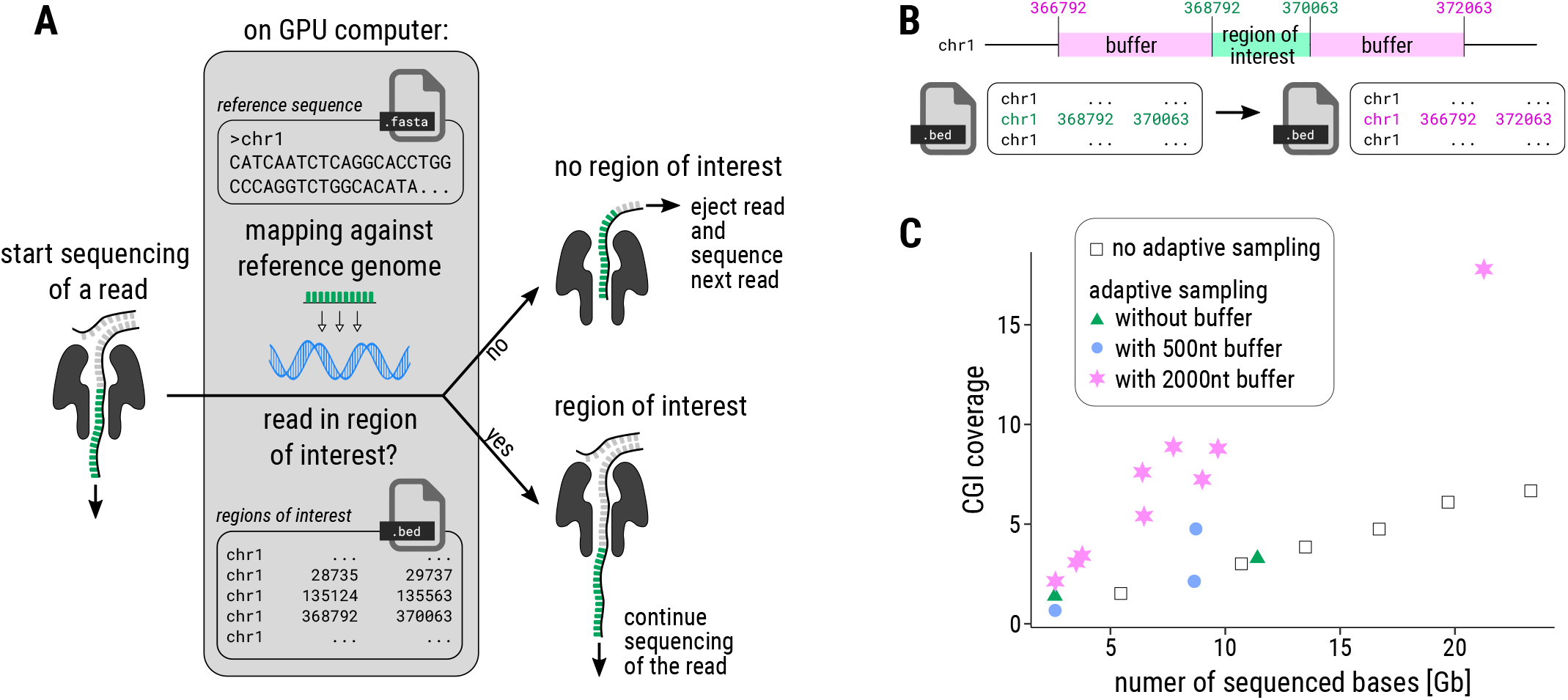
**(A)** The basic principle of adaptive sampling: For each read, the first ∼400 bases are mapped to a given reference, and sequencing continues only if they match a region of interest (22). About 450 bases are sequenced per second for R9nanopore flow cells (2). Rejecting a read takes about 0.5 seconds, with additional time required to capture the next read. **(B)** In order not to miss the reads where the region of interest (green region) is not at the beginning of the read, the target region should be defined with an extended buffer (e.g. 2000 nt) on both sides (pink regions) in the configuration .bed file. **(C)** Adaptive sampling was used to enrich over 30,000 CpG islands (CGIs) to increase their coverage. To increase enrichment, each CGI region of interested was extended by a 500 nt buffer region (blue dots) and by a 2000 nt buffer region (pink stars) on both sides.

As the acceptance or rejection of a read is based solely on its first chunk, strands may be rejected, if the region of interest is not located at the beginning of the read. To address this, the target region of interest should be extended by adding a “buffer” on both sites in the configuration file to accommodate sequencing in either direction, as sequencing can initiate from either DNA strand (Fig. 8B). Thus, for adaptive sampling, buffer describes an amount of nucleotides, by which the region of interest is extended in the `bed` file (a text file format used to store genomic regions).

Using adaptive sampling may yield a higher number of short reads (reducing N50 and median read length), and necessitating re-basecalling with a higher accuracy model post-sequencing to enhance data quality.

We have applied adaptive sampling to enrich CpG islands in the human genome, see Tab. S5. We targeted approx. 30 000 CpG islands, with an average length of 777 nt (IDs 153.1– 178.1). We applied adaptive sampling without adding a buffer region, with a 500 nt buffer region, and a 2 000 nt buffer region on both sides of the target regions. We achieved up to 17 X mean sequencing depth on CpG islands, representing a fourfold increase compared to the mean sequencing depth of the whole genome, when applying 2 000 nt buffer. According to ONT, the theoretical maximum increase in sequencing depth achievable through adaptive sampling is approximately 5–10-fold^15^. In summary, adaptive sampling can effectively enhance sequencing depth in specific genomic regions or target species, albeit with increased time requirements for post-processing, such as re-basecalling and filtering out short, unwanted reads.

### CHALLENGES IN ANALYZING ONT DATA

#### Assembling an ONT pipeline

Bioinformatics analysis of ONT sequencing data is based on modular pipelines, that can be assembled and utilized effectively, even by users with limited computational backgrounds (Fig. 9). Instead of relying solely on standard, one-size-fits-all pipelines, it is critical to carefully select tools that best align with specific experimental goals and data characteristics.

**Figure 9.**
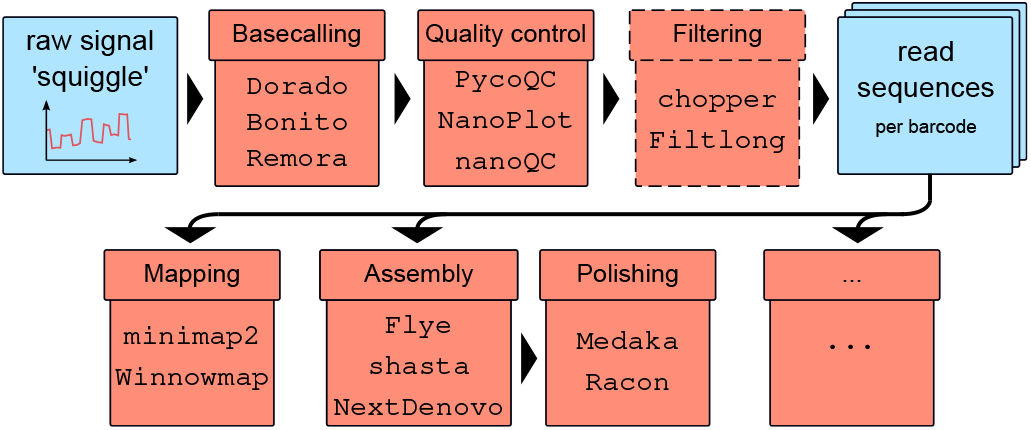
The standard pipeline for nanopore sequencing data involves basecalling of the raw data, followed by quality control and filtering. The reads can then be analyzed with specialized long-read tools for mapping, polished assembly, or other workflows.

A well-constructed ONT analysis pipeline includes several steps, which can be addressed by different tools:

**Basecalling** is the transformation of raw signal data, or ‘squiggles,’ into the original nucleotide sequences using neural networks. Typically, the basecalling tools have a pre-processing step, such as removing the adaptors and normalizing individual signals. The following three tools are all provided by ONT. Currently, `Dorado` ^16^ is the state-of-the-art software for basecalling on ONT platforms, that supports the older `fast5` and the new `pod5` format, which is a more compact format enabling faster and more efficient file access. `Dorado` includes basecalling models for all ONT kits, predicts modified bases, and built-in demultiplexing capabilities for barcode detection from ONT barcoding kits. `Bonito` ^17^ is an alternative tool that allows users to train their own neural networks for basecalling. `Remora` ^18^ adds an additional layer, enabling nucleotide modification calling by training neural networks along the standard bases. Alternative tools for nucleotide modifications prediction are discussed in detail below. Basecalling results are typically saved in `fastq` format for subsequent analysis steps.

After basecalling, **quality control** (QC) is essential to evaluate the reliability of the sequencing data and to identify any potential issues in the dataset, such as degradation, contamination, and read length distribution. One of the most comprehensive QC tools for ONT data is `pycoQC` ^19^ (64), which provides an interactive `html` overview that includes information on read quality, read length, sequence coverage, active channels, and quality over sequencing time. Other QC tools, such as `NanoPlot` ^20^ (65) and `nanoQC` ^21^, provide a restricted functionality, but with a different representation of the data.

Following QC, **filtering** is often necessary, especially for experiments requiring specific read quality or length thresholds. Filtering tools help to remove low-quality reads, adapters, barcodes, or reads outside the desired range, increasing the overall quality and relevance of the dataset. `chopper` ^22^ from the `NanoPack` suite (65) or `Filtlong` ^23^ are widely used for filtering ONT reads. `Dorado` also provides parameters to trim adapters and barcodes.

Once high-quality reads are obtained, subsequent analyses such as mapping and assembly can be performed using specialized tools designed for long-read data. For read **mapping**, `minimap2` ^24^ (66) is widely used. `Winnowmap` ^25^ (67) is an alternative mapping tool that has been specifically optimized for complex genomes. It has demonstrated superior performance when mapping both simulated PacBio and ONT data for human genomes (67). For assembly, `Flye` is one of the most widely used tools, demonstrating good performance in terms of contig length and error rate in bacteria (68). It includes polishing and can be told how many iterations of polishing should be performed. Alternative tools, such as `Shasta` ^26^ (69), have been shown to perform faster, while `NextDenovo` ^27^ (70) can generate longer contigs, particularly in the case of mollusks (71). **Error correction** (polishing) can be subsequently utilized to refine the assembly. `Medaka` ^28^ is a ONT-specific neural networks, trained on typical ONT errors to correct these errors effectively. `Medaka` has proven particularly effective in reducing deletion errors (68). Additionally, `Racon` ^29^ (72) is a consensus caller using high quality reads from technologies such as Illumina to reduce single nucleotide variants and insertion errors, enhancing the overall accuracy of the assembled data.

#### Hidden treassures in raw data

Beyond the standard pipeline of ONT tools, the raw data from ONT sequencing contains a hidden treasure of information. Analyzing this data requires significant self-implementation and detailed knowledge. Here, a demonstration of the fundamentals necessary to fully exploit the potential of ONT by using DNA and RNA modification detection as an example is given. This description will pave the way to complement existing tools for nucleotide modification analysis, which are currently designed for only a small fraction of modifications. To achieve this, we will focus on processing the raw data, which can be divided into five distinct phases: (1) Accessing raw data: in this initial step, the raw signal data from the nanopore sequencer is imported into the computational analysis. This is particularly important because it serves as the foundation for all subsequent analyses. (2) Improving Signal Segmentation: is applied to refine the process of identifying individual signals corresponding to nucleotides. Improving segmentation helps enhance the accuracy of the subsequent basecalling step by ensuring that the software can effectively distinguish between different signal components. (3) DNA Modification Detection: Once the signal data is well-segmented, this step focuses on identifying specific modifications to DNA (e.g., methylation). Detection of DNA modifications is important for understanding epigenetic regulation and gene expression, providing insights into cellular functions and biological processes. (4) RNA Modification Detection: Similar to DNA modification detection, this step involves identifying modifications in RNA sequences. RNA modifications can influence gene expression and stability, making their detection crucial for studying transcriptomics and understanding post-transcriptional regulation. (5) Differential Signal Detection: Alternatively to predicting modification types directly, the differences in signal patterns between conditions or samples can be analyzed to reveal a general change of the nucleotide. **Preprocessing raw data**. ONT stores the ‘squiggles’ as integers (73) to save storage space, which are converted back to pA, see SEqu. **1**. Normalization of raw ONT signals is essential for reliable comparisons within and across sequencing experiments. Signal variations caused by differences in flow cells, nanopores, and sensors can introduce biases (Fig. S11), making normalization (Fig. 10A) critical for accurate and reproducible analyses (SEqu. **2**). Outliers in ONT signals, as in Fig. 10B, often caused by faulty sensor measurements, can distort downstream analyses. To ensure data quality, these outliers should be identified and filtered prior to further signal processing. The ONT signal, comprising a continuous series of data points, may occasionally include multiple adjacent sequencing reads, as shown in Fig. 10C.

**Figure 10.**
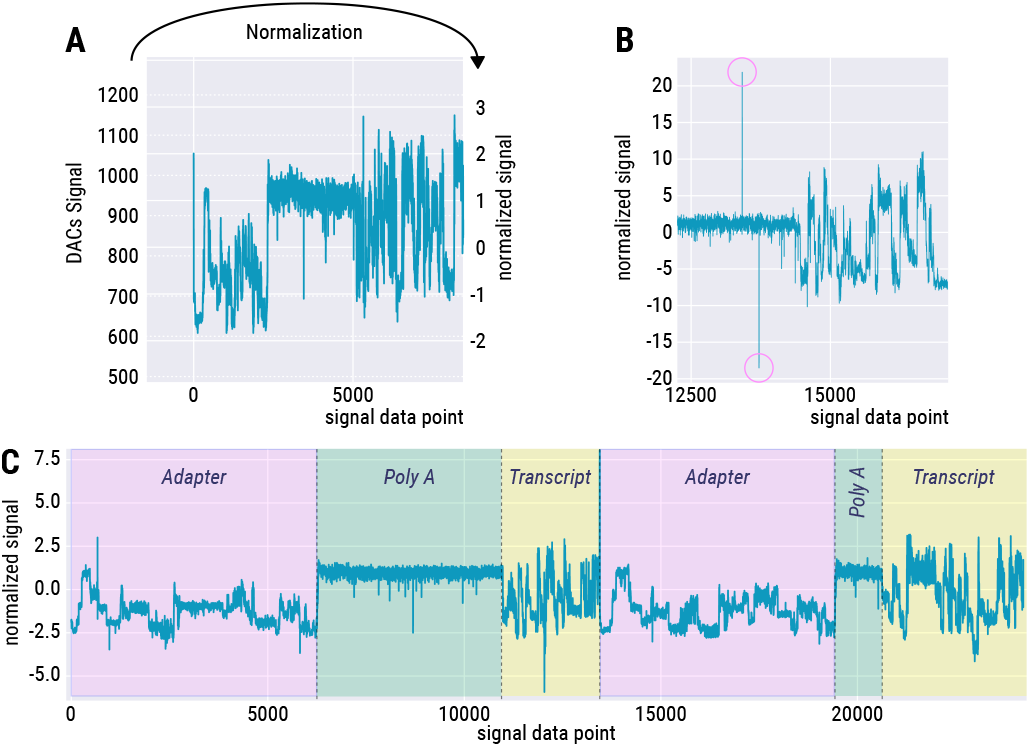
Overview of Signal Data Preprocessing Steps. **A** To standardize the data, the unnormalized time series (left) are converted (arrow) into the corresponding normalized signal (right, mostly ranging from -4 to 4). **B** Detected outliers (pink circles) indicate points that deviate significantly from the expected signal pattern, likely due to sensor errors. **C** A single signal can contain more than one read. Here indicated with signal containing three recurring components for ‘Read 1’ and ‘Read 2’ specific to RNA sequencing: ‘adapter,’ ‘poly A,’ and ‘transcript.’

The reason for this phenomenon is unresolved. To ensure accurate and reliable downstream analyses, such signals must be split into individual reads using the coordinates provided by `Dorado`.

The python package `read5` ^30^ is a wrapper that unifies access to `fast5`, `pod5`, and `slow5` data formats and provides functions for signal conversion (SEqu. **1**) and normalization (SEqu. **2**). Each of these formats has its own API using different function names which often requires to script format-specific code to handle inconsistencies.

##### Improving signal segmentation

The analysis of raw data plays a crucial role for detecting chemical nucleotide modifications. This is particularly relevant for the vast amount of around 170 RNA modifications for which no basecalling models currently exist. These modifications also produce characteristic signals when translocating through the nanopore. Analyzing raw signals for modification detection requires improved signal segmentation, known as ‘resquiggling.’ This process involves realigning the raw current signal to the nucleotides, see Fig. 11. This segmentation is crucial because the motor protein that moves the nucleotides through the pore does not operate at a constant speed. The time a nucleotide spends in the pore (‘dwell time’) varies depending on the specific nucleotides (74).

**Figure 11.**
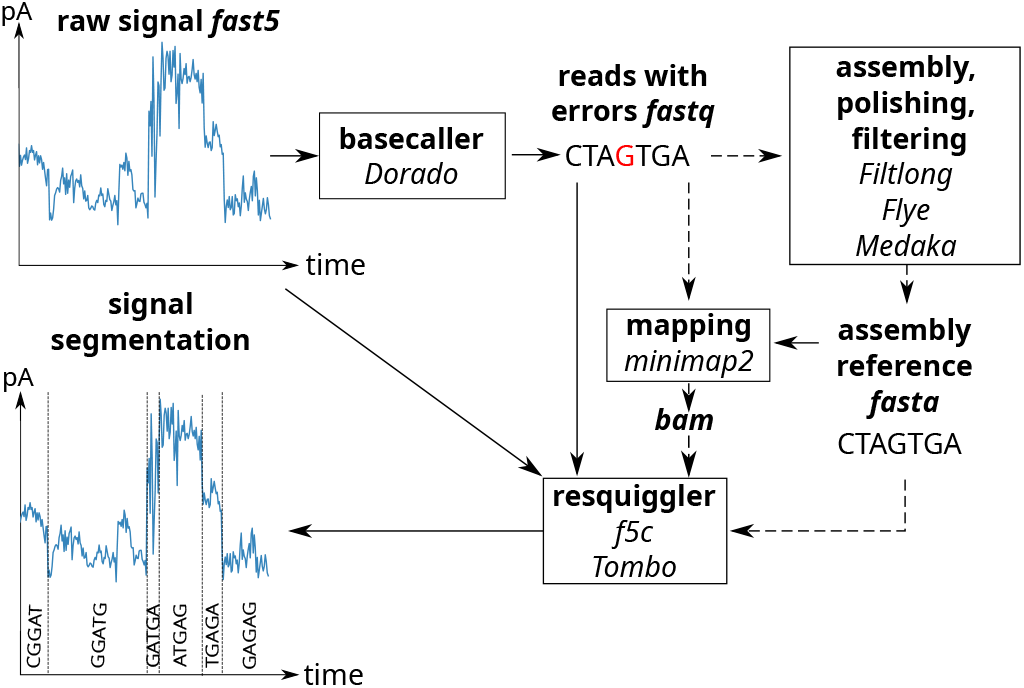
Overview of Signal Processing and Modification Calling. The raw ONT signal can be segmented and corrected for basecalling errors by resquigglers using the raw signal and basecalls. Some resquiggling tools also require a reference sequence and a mapping.

Several tools have been developed to facilitate the resquiggling process and require generally (i) the raw ONT signal (`fast5`, `slow5` or `pod5`) and (ii) the basecalls generated by the basecalling process (`fastq`). Optionally, (iii) a reference sequence (`fasta`) and (iv) the alignments of the basecalls to the reference (`bam`) can be added. The most widely used tool `f5c` (75) is a GPU-accelerated, multi-threaded re-implementation of `Nanopolish Eventalign`. It offers significant reductions in runtime while maintaining the known output formats. Notably, `f5c` also supports newer pore chemistries, such as R10 and RP4. For instance, a sequencing run of a 32,000 bp Coronavirus genome (ID 13.1) with 1.3 Gb across 857,000 reads produced a 181.5 GB output file using `Nanopolish Eventalign`, which has been reduced to 84 GB with `f5c`. `Tombo` has not been supported by ONT since 2020 and is compatible only with older single-read `fast5` formats. Despite its limitations, `Tombo` remains in use today for specific applications.

Future developments in resquiggling tools are expected to further enhance compatibility with emerging nanopore technologies and data formats.

##### Detecting DNA modifications

The direct detection of DNA modifications without amplification or pre-treatment of the DNA is an exciting feature of nanopore sequencing, providing insights into epigenetic regulation and genomic variability. DNA modifications alter the electrical signal as the DNA passes through the nanopore resulting in a differing picoampere signal for k-mers (for R9 k=5; for R10 k=9) containing modified bases compared to those consisting of only unmodified bases, see Fig. 12. Currently, 17 types of DNA modifications have been identified in the genomes of bacteria and eukaryotes (76), with the most well-characterized being methylation of cytosine 5-methylcytosine, 5mC (77); 5-hydroxymethylcytosine, 5hmC; and adenine N6-methyladenine, 6mA (78).

**Figure 12.**
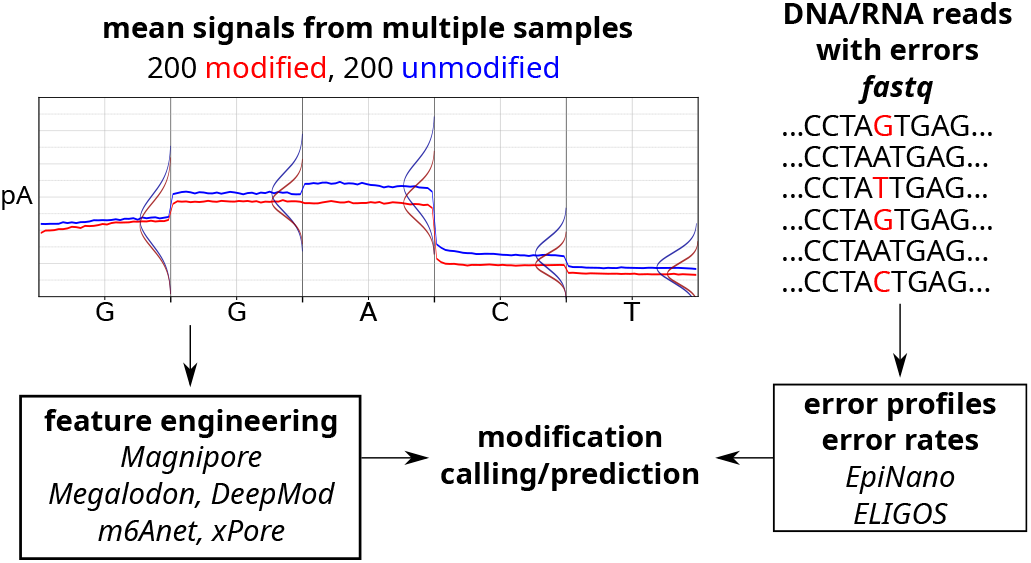
Modified nucleotides have their own characteristic signal. Machine learning models can detect this signal directly or by comparing it to the unmodified counterpart. The red and blue lines represent the mean signal of 200 reads respectively, while the vertical bulges indicate the signal distribution per base following a normal distribution. Modifications can lead to error patterns in the reads that can be detected by tools.

Several computational tools have been developed to identify modifications from nanopore data, with a primary focus on 5mC and 6mA, using different computational approaches, in particular Hidden Markov Models, statistical tests, or neural networks, see Tab. S2. Input data for the methylation calling tools is usually the raw or basecalled `fast5` /`pod5` files.

DNA modification calling was traditionally performed after basecalling using tools like `Nanopolish` (79) or ONT’s `Megalodon` ^31^. Currently, ONT’s `Dorado` ^32^ detects CpG methylations directly during basecalling if specified, and provides greater accuracy than tools like `Nanopolish` (80), which has difficulty when nearby CpG sites differ in methylation (79). The performance of modification-detection tools varies by slight over-prediction (e.g. `Nanopolish`) or under-prediction (e.g. `DeepMod`) as shown in several reviews (21, 80, 81, 82, 83) Validation remains essential, although tools like `Nanopolish`, `Megalodon`, `DeepSignal`, and `Dorado` already show high correlation with the gold standard, whole-genome bisulfite sequencing (WGBS) (80, 81). Meta approaches improve modification detection accuracy by combining the outputs of multiple tools. For example `METEORE` combines results from up to six tools using a random forest model, significantly improving accuracy but at the cost of increased runtime (21).

##### Detecting RNA modifications

The detection of RNA modifications using nanopore sequencing is less developed compared to DNA modifications, possibly because RNA is known to contain over 170 distinct modifications (84). RNA modifications are of utmost importance for RNA function and regulation.

Computational tools exist only for a handful of RNA modifications, see Tab. S2. Beside these downstream analysis tools, ONT’s `Dorado` directly enables modification detection by specifying specific basecalling models. Users can further customize the process by manually disabling additional filters as needed. Excitingly, basecallers like `Bonito` and `Remora`, allow researchers to train custom models for detection of modifications of any kind. These models can be created from scratch or fine-tuned from pre-trained versions, offering flexibility for different datasets and research objectives. The models can perform binary classification to predict a certain modification, or directly call modified bases alongside the standard nucleotide sequence.

##### Detecting differential signals

Comparing raw ONT signals between different samples provides a powerful approach for detecting differential signals, which can point to mutations, modifications and isotopic labeling (85). The goal of this approach is not necessarily to identify the exact type of modification but rather to detect the presence of a modification. The signal may differ in its measured pA range, including mean, variance, skewness, kurtosis, and dwell time. The calculation of these measures can be applied to full-length nanopore reads. To minimize normalization biases, comparative analyses should ideally be performed using the same flow cell. When comparisons across multiple flow cells are unavoidable, further normalization steps are needed.

On the other hand, for example, in the case of modifications such as deuterium in the DNA/RNA backbone (85), normalization can inadvertently obscure biologically meaningful information. This is particularly true when the signal-to-noise ratio—where the signal represents the modification and the noise stems from instrumental bias—is low. To address these challenges, carefully controlled experimental designs are essential to ensure that observed differences in the nanopore signal reflect genuine modifications rather than artifacts of the sequencing process.

## CONCLUSION

In summary, this study has yielded critical insights into refining nanopore sequencing using the MinION platform, with far-reaching implications for research and application. We displayed suggestions for the user of nanopore sequencing in the three categories (1) library preparation tips; (2) technical tips; and (3) analyzing ONT data, see a summary in Fig. 13. We showed general statistics for the number of sequenced bases after 12 h on both R9 and R10 flow cells. EB12 is influenced by the number of pores and channels at the start of the run for DNA, as well as the pore half time, which surprisingly varies depending on the sample type. In contrast, EB12 shows no correlation with the amount of input material, read length, or flow cell age. Additionally, we showed, that the flow cell half-life depends for DNA samples on the buffer used (LFB vs. SFB) and that flow cells can perform well after they have been expired. Meanwhile, we could also verify most of the observations on R10.4 flow cells. However, for the current time point the sample size is too small to make statistically valid statements.

**Figure 13.**
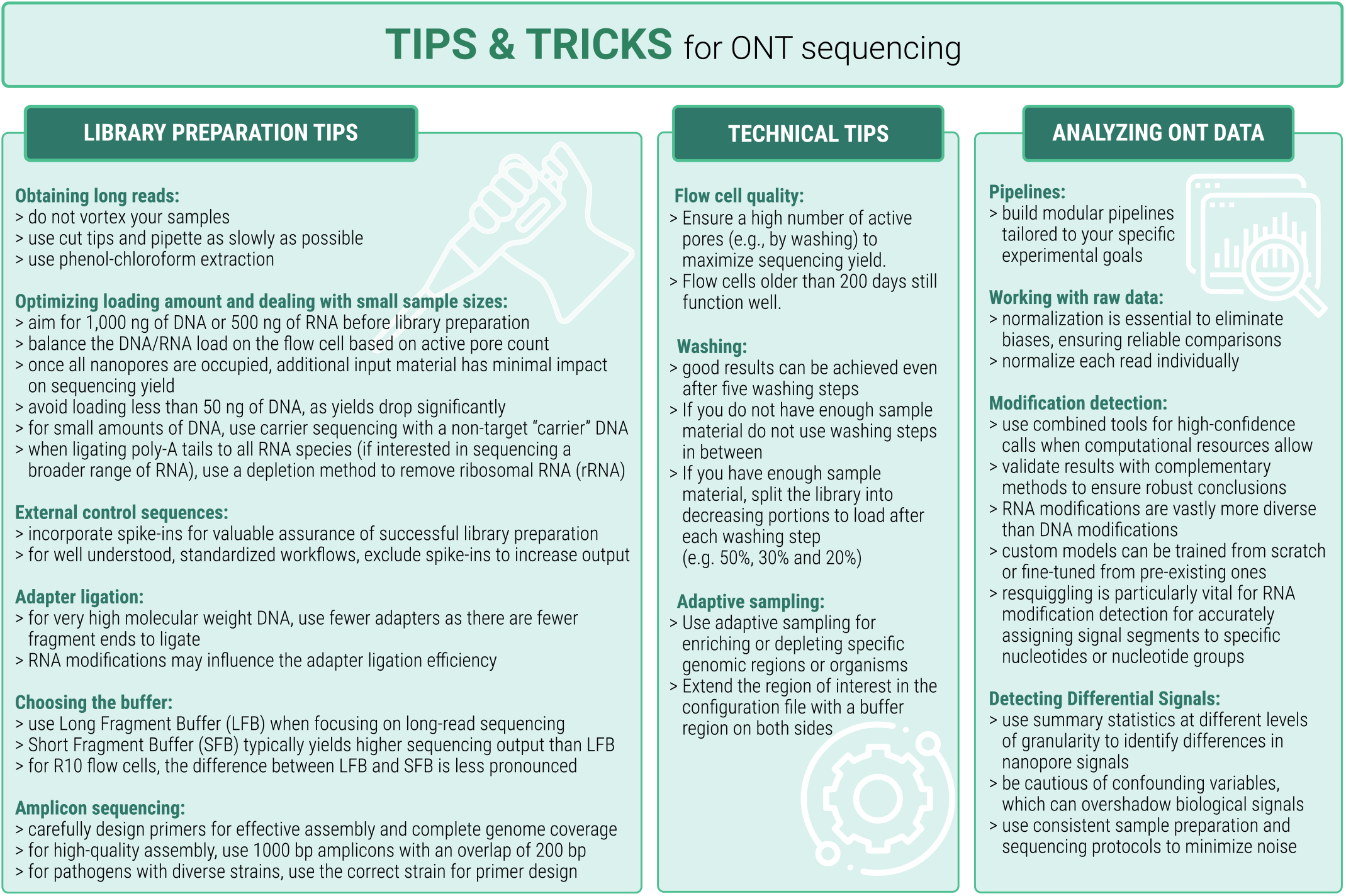
TIPS & TRICKS mentioned within the manuscript are summarized here for a better overview.

With this publication, correlations and assertions from the community have been statistically substantiated and are now citable for future work.

Here, we aim to describe an additional observation: within the community, it is often posited claimed that theoretically, one could sequence reads of unlimited length if the DNA fragments were arbitrarily long. We extracted fragments of 500,000 nt from algae *Chlamydomonas euryale, Chlamydomonas reinhardtii*; however, we encountered significant challenges during the sequencing process. Despite trying various methods, we consistently observed good sequencing performance only in the initial minutes before the sequencing had to be aborted. Regarding this observation, we currently have a few potential explanations: (1) the molarity of the adaptors with respect to the high molecular weight fragments might have been incorrect; or (2) algae may contain a high amount of polysaccharides that could influence pore activity. As seen for this example many of the observations can still not be entirely explained and call systematic analysis.

## ACKNOWLEDGEMENTS

We thank Franziska Hufsky for her valuable assistance in improving the writing of this manuscript and Emanuel Barth for his valuable input on statistical questions. We thank Carolin Dippmann for her experimental work and insightful discussions that contributed to the development of this study and Franziska Aron for her support and help in the lab. We thank Marie Lataretu for her contribution to the ONT DNA spike-in and Adrian Viehweger for his work on figures. We thank Patrick Bohn for discussion about different flow cell types. We thank the Sequencing Core Facility of the Genome Competence Center (MF1) of the Robert Koch Institute (RKI), Germany for providing excellent sequencing services and André Götze from the RKI MF1 DigiLab for outstanding support in obtaining technical metadata information of the associated runs. We thank Matthew Huska and Christian Blumenscheit from MF1 for R10.4.1 data obtained in the context of an RKI special research funds project about adaptive sequencing of a Zymo Mock community. Competing interests: DM and BS are employees of oncgnostics GmbH, a company that aims to commercialize DNA methylation markers. This work was funded by the Deutsche Forschungsgemeinschaft (DFG, German Research Foundation) under Germany’s Excellence Strategy – EXC 2051 – Project-ID 390713860, the German DFG Collaborative Research Centre AquaDiva (CRC 1076 AquaDiva), the German state of Thuringia via the Thüringer Aufbaubank (2021 FGI 0009), the Carl-Zeiss-Stiftung within the program Scientific Breakthroughs in Artificial Intelligence (project “Interactive Inference”), and the Ministry for Economics, Sciences and Digital Society of Thuringia (TMWWDG), under the framework of the Landesprogramm ProDigital (DigLeben-5575/10-9).

## SUPPLEMENTARY INFORMATION

**Table S1:** Sample Summary. overview of all runs used for the analysis of Fig. 2, Fig. 4, Fig. 6 and Fig. S4–S10. RNA or DNA sequencing; fc version – R9 or R10 flow cells; device – MinION or GridION device; #cha – amount of active channels at the begin of the sequencing run; #por – amount of active pores at the beginning of the sequencing run; fc age – time passed since delivery of the Flow cell in days; sample type – from which kind of species the sequenced material originated in nine categories: virus, bacteria, protist, metagenomic, insect, mouse, human; buffer – LFB, SFB or RNA buffer; ng load – amount of loaded library (DNA/RNA) in *ng*; EB 12 – amount of sequenced bases after 12 h, estimated by sequencing software MinKNOW; mean rl – mean estimated read length of all sequenced reads in bases; pore hl – calculated pore half life time of the flow cell in hours (see main text).

| ID | RNA/<br>DNA | fc<br>version | device | #cha | #por | fc<br>age | sample<br>type | buffer | ng<br>load | Estimated<br>bases | EB12 | mean<br>rl | pore<br>hl | fc<br>id |
| --- | --- | --- | --- | --- | --- | --- | --- | --- | --- | --- | --- | --- | --- | --- |
| 1.1 | DNA | R10 | GridIon | 510 | NA | NA | virus | SFB | NA | 8161 | 4721 | 2036 | 998 | FAU91805 |
| 2.1 | DNA | R10 | GridIon | 480 | NA | NA | virus | SFB | NA | 4745 | 4745 | 2934 | 1630 | FAX97604 |
| 3.1 | DNA | R10 | GridIon | 467 | 1766 | NA | virus | SFB | NA | 6041 | 3977 | 2137 | 643 | FAY00760 |
| 4.1 | DNA | R10 | GridIon | 502 | 1860 | NA | virus | SFB | NA | 8139 | 5074 | 2640 | 1012 | FAY01081 |
| 5.1 | DNA | R10 | GridIon | 501 | 1869 | NA | virus | SFB | NA | 9339 | 5444 | 3776 | 1397 | FAY01249 |
| 6.1 | DNA | R10 | GridIon | 504 | NA | NA | virus | SFB | NA | 9932 | 5699 | 4022 | 1510 | FAY01385 |
| 7.1 | DNA | R10 | MinIon | 293 | 1368 | 172 | virus | SFB | 60 | 177 | 153 | 419 | 51 | FAO88286 |
| 8.1 | RNA | R9 | MinIon | 295 | 749 | 174 | virus | RNA | 140 | 523 | 423 | 1530 | 900 | FAM95379 |
| 9.1 | RNA | R9 | MinIon | 487 | 1333 | 56 | virus | RNA | 210 | 1571 | 923 | 1700 | NA | FAO07649 |
| 10.2 | RNA | R9 | MinIon | 489 | 1385 | 49 | virus | RNA | 90 | 1495 | 1031 | 1454 | NA | FAO10418 |
| 11.1 | RNA | R9 | MinIon | 501 | 1282 | 10 | virus | RNA | 74 | 1324 | 591 | 1072 | 1638 | FAO53185 |
| 12.1 | RNA | R9 | MinIon | 506 | 1372 | 112 | virus | RNA | 308 | 1247 | 885 | 1046 | 797 | FAO83518 |
| 13.1 | RNA | R9 | MinIon | 473 | 1201 | 127 | virus | RNA | 186 | 1326 | 869 | 1577 | 810 | FAO86549 |
| 14.1 | RNA | R9 | MinIon | 470 | 1201 | 102 | virus | RNA | 304 | 118 | 117 | 1206 | 79 | FAO86555 |
| 15.1 | RNA | R9 | MinIon | 505 | 1380 | 111 | virus | RNA | 392 | 1028 | 801 | 1123 | 637 | FAO86826 |
| 16.1 | RNA | R9 | MinIon | 510 | 1342 | 34 | virus | RNA | 256 | 483 | 462 | 982 | 269 | FAO86858 |
| 17.1 | RNA | R9 | MinIon | 500 | 1241 | 111 | virus | RNA | 348 | 136 | 136 | 1112 | 113 | FAO86895 |
| 18.1 | RNA | R9 | MinIon | 500 | 1308 | 110 | virus | RNA | 266 | 626 | 580 | 1099 | 414 | FAO86907 |
| 19.1 | RNA | R9 | MinIon | 498 | 1298 | 110 | virus | RNA | 420 | 296 | 296 | 1140 | 212 | FAO86991 |
| 20.1 | RNA | R9 | MinIon | 442 | 1142 | 93 | virus | RNA | 224 | 284 | 283 | 1354 | 225 | FAO86996 |
| 21.1 | RNA | R9 | MinIon | 495 | 1228 | 111 | virus | RNA | 360 | 314 | 312 | 1133 | 215 | FAO87031 |
| 22.1 | RNA | R9 | MinIon | 482 | 1249 | 127 | virus | RNA | 276 | 295 | 294 | 1353 | 232 | FAO87106 |
| 23.1 | RNA | R9 | MinIon | 432 | 1002 | 9 | virus | RNA | 167 | 1400 | 650 | 1677 | 1348 | FAO88388 |
| 24.1 | DNA | R9 | MinIon | 512 | 1960 | 3 | virus | SFB | 575 | 13251 | 6708 | 765 | 1439 | FAO92019 |
| 25.1 | RNA | R9 | MinIon | 491 | 1327 | 110 | virus | RNA | 346 | 600 | 556 | 1111 | 352 | FAO92042 |
| 26.1 | DNA | R9 | MinIon | 511 | 1907 | 6 | virus | SFB | 516 | 9748 | 6396 | 866 | 958 | FAO92112 |
| 27.1 | RNA | R9 | MinIon | 497 | 1307 | 110 | virus | RNA | 356 | 768 | 657 | 1123 | 526 | FAO92116 |
| 28.1 | RNA | R9 | MinIon | 502 | 1365 | 111 | virus | RNA | 254 | 1141 | 847 | 975 | 795 | FAO92254 |
| 29.1 | RNA | R9 | MinIon | 322 | 893 | 93 | virus | RNA | 236 | 317 | 316 | 1151 | 368 | FAO92303 |
| 30.1 | RNA | R9 | MinIon | 350 | 1210 | 167 | virus | RNA | 268 | 1708 | 767 | 1666 | 1373 | FAP02359 |
| 31.1 | RNA | R9 | MinIon | 93 | 152 | 51 | virus | RNA | 580 | 46 | 44 | 1256 | 166 | FAQ01609 |
| 31.2 | RNA | R9 | MinIon | NA | 814 | NA | virus | RNA | 580 | NA | 487 | 1246 | 557 | FAQ01609 |
| 32.1 | RNA | R9 | MinIon | 420 | 965 | 147 | virus | RNA | 52 | 7 | 7 | 605 | 53 | FAQ69187 |
| 33.1 | DNA | R10 | GridIon | 510 | 1609 | NA | bacteria | LFB | NA | 22320 | 4567 | 10233 | 3562 | FAU91295 |
| 34.1 | DNA | R10 | GridIon | 511 | 1556 | NA | bacteria | LFB | NA | 10173 | 3988 | 7108 | 2071 | FAU92937 |
| 35.1 | DNA | R10 | GridIon | 502 | 1491 | NA | bacteria | LFB | NA | 12166 | 3944 | 7911 | 1930 | FAW21704 |
| 36.1 | DNA | R10 | GridIon | 510 | 1460 | NA | bacteria | LFB | NA | 12767 | 3779 | 1553 | 2366 | FAW22654 |
| 37.1 | DNA | R10 | GridIon | 510 | 1578 | NA | bacteria | LFB | NA | 14943 | 4433 | 3423 | 2026 | FAW22671 |
| 38.1 | DNA | R10 | GridIon | 348 | 1019 | NA | bacteria | LFB | NA | 6458 | 2713 | 1629 | 1675 | FAW23628 |
| 39.1 | DNA | R10 | GridIon | 498 | 1567 | NA | bacteria | LFB | NA | 6065 | 3604 | 6144 | 719 | FAW45145 |
| 40.1 | DNA | R10 | GridIon | 510 | 1566 | NA | bacteria | LFB | NA | 14424 | 4395 | 8707 | 2285 | FAW45147 |
| 41.1 | DNA | R10 | GridIon | 478 | 1501 | NA | bacteria | LFB | NA | 7288 | 3437 | 9545 | 1050 | FAW45167 |
| 42.1 | DNA | R10 | GridIon | 413 | 1262 | NA | bacteria | LFB | NA | 2715 | 2177 | 1855 | 456 | FAW98004 |
| 43.1 | DNA | R10 | GridIon | 503 | NA | NA | bacteria | LFB | NA | 3563 | NA | NA | NA | FAW98056 |
| 44.1 | DNA | R10 | GridIon | 499 | 1899 | NA | bacteria | LFB | NA | 13610 | 6244 | 3383 | 1169 | FAW99941 |
| 45.1 | DNA | R10 | GridIon | 495 | NA | NA | bacteria | LFB | NA | 4855 | NA | NA | NA | FAX00084 |
| 46.1 | DNA | R10 | GridIon | 484 | 1484 | NA | bacteria | LFB | NA | 7281 | 3929 | 2441 | 851 | FAX00224 |
| 47.1 | DNA | R10 | GridIon | 487 | 1812 | 105 | bacteria | LFB | 1216 | 11256 | 4949 | 3599 | 934 | FAX12900 |
| 48.1 | DNA | R10 | GridIon | 221 | 813 | 112 | bacteria | LFB | 403 | 4209 | 2002 | 3186 | 657 | FAX13587 |
| 49.1 | DNA | R10 | GridIon | 511 | NA | 98 | bacteria | LFB | 552 | 14912 | 5538 | 1931 | NA | FAX14134 |
| 50.1 | DNA | R10 | GridIon | 506 | NA | 105 | bacteria | LFB | 324 | 10673 | 4891 | 2555 | NA | FAX18203 |
| 51.1 | DNA | R10 | GridIon | 484 | 1844 | NA | bacteria | LFB | NA | 21675 | 5881 | 3121 | 2151 | FAX70780 |
| 52.1 | DNA | R10 | MinIon | 235 | 999 | 186 | bacteria | SFB | 139.2 | 1239 | 770 | 1169 | 1531 | FAO85473 |
| 53.1 | DNA | R10 | MinIon | 348 | NA | 36 | bacteria | LFB | NA | 2625 | NA | NA | NA | FAS37108 |
| xx.1 | DNA | R10 | MinIon | 428 | 1116 | 35 | bacteria | LFB | 1008 | 1242 | 1041 | 6008 | NA | FAS37108 |
| 54.1 | DNA | R10 | MinIon | 132 | NA | 31 | bacteria | LFB | 298 | 997 | 707 | 9448 | NA | FAS38416 |
| xx.1 | DNA | R10 | MinIon | 417 | NA | 30 | bacteria | LFB | 276 | 4825 | 3281 | 9328 | NA | FAS38416 |
| 55.1 | DNA | R10 | MinIon | 229 | NA | 1 | bacteria | LFB | 1440 | 651 | 443 | 2796 | NA | FAV74660 |
| 56.1 | DNA | R10 | MinIon | 500 | NA | 7 | bacteria | LFB | 157 | 11585 | 4816 | 3574 | NA | FAW66139 |
| 57.1 | DNA | R10 | MinIon | 468 | NA | 2 | bacteria | LFB | NA | 9767 | 4480 | 5356 | NA | FAW73005 |
| xx.1 | DNA | R10 | MinIon | NA | NA | 1 | bacteria | LFB | NA | NA | NA | NA | NA | FAW73005 |
| 58.1 | DNA | R10 | MinIon | 226 | NA | 1 | bacteria | LFB | 199 | 4678 | NA | NA | NA | FAW75095 |
| 59.1 | DNA | R10 | MinIon | 487 | NA | 1 | bacteria | LFB | 226 | 0 | NA | NA | NA | FAX18284 |
| 60.1 | DNA | R10 | MinIon | 473 | NA | 1 | bacteria | LFB | 154 | 12573 | 4714 | 4304 | NA | FAX30985 |
| 61.1 | DNA | R9 | GridIon | 484 | 1447 | NA | bacteria | LFB | NA | 9421 | 3623 | 7487 | 1791 | FAU91863 |
| 62.1 | DNA | R9 | GridIon | 511 | 1398 | NA | bacteria | LFB | NA | 6066 | 3190 | 13070 | 2225 | FAV24719 |
| 63.1 | DNA | R9 | GridIon | 494 | 1512 | NA | bacteria | LFB | NA | 18854 | 6177 | 9601 | 2187 | FAV30532 |
| 64.1 | DNA | R9 | MinIon | 139 | 482 | 101 | bacteria | LFB | 200 | 100 | NA | 5093 | 18 | FAO16410 |
| 65.1 | DNA | R9 | MinIon | 499 | 1439 | 4 | bacteria | SFB | 238 | 18063 | 6136 | 6785 | 2038 | FAO53260 |
| 66.1 | DNA | R9 | MinIon | 416 | 1021 | 114 | bacteria | SFB | 352 | 11310 | 4237 | 3302 | 2384 | FAO87074 |
| 67.4 | DNA | R9 | MinIon | 449 | 1541 | 37 | bacteria | LFB | 200 | 654 | NA | 4932 | NA | FAO92001 |
| 68.1 | DNA | R9 | MinIon | 416 | 1161 | 178 | bacteria | SFB | 600 | 7394 | 4172 | 2023 | 1108 | FAP00827 |
| 69.1 | DNA | R9 | MinIon | 449 | 1659 | 121 | bacteria | SFB | 775 | 2507 | NA | 18442 | NA | FAQ01004 |
| 69.2 | DNA | R9 | MinIon | 355 | 1393 | 122 | bacteria | SFB | 444 | 5911 | 3731 | 12254 | 2 | FAQ01004 |
| 70.1 | DNA | R9 | MinIon | 494 | 1620 | 44 | bacteria | SFB | 1428 | 8812 | 4452 | 1262 | 1652 | FAQ58659 |
| 71.1 | DNA | R9 | MinIon | 454 | 1692 | 58 | bacteria | SFB | 504 | 18406 | 5950 | 20443 | 2303 | FAQ58677 |
| 72.1 | DNA | R9 | MinIon | 493 | 1312 | 41 | bacteria | SFB | 479.6 | 25403 | 6853 | 2023 | 2787 | FAQ69194 |
| 73.1 | DNA | R9 | MinIon | 480 | 1468 | 87 | bacteria | LFB | 700 | 23019 | 6807 | 23746 | 2554 | FAR96927 |
| 74.1 | DNA | R9 | MinIon | 406 | 1398 | 14 | protist | LFB | 600 | 4265 | 3028 | 1592 | 446 | FAQ01569 |
| 75.1 | DNA | R9 | MinIon | 485 | 1692 | 80 | protist | SFB | 310 | 4620 | 3452 | 1988 | 608 | FAQ01665 |
| 76.1 | DNA | R9 | MinIon | 508 | 1881 | 16 | protist | SFB | 528 | 10925 | 5862 | 1852 | 1169 | FAQ01773 |
| 77.1 | DNA | R9 | MinIon | 499 | 1785 | 78 | protist | SFB | 470 | 4948 | 3880 | 1867 | 557 | FAQ01847 |
| 77.2 | DNA | R9 | MinIon | 371 | 1329 | 79 | protist | SFB | 160 | 2085 | 1749 | 2093 | 258 | FAQ01847 |
| 78.1 | DNA | R10 | GridIon | 503 | 1719 | NA | metagenomic | LFB | NA | 5046 | 2434 | 5637 | 821 | FAW10206 |
| 79.1 | DNA | R10 | GridIon | 501 | 1723 | NA | metagenomic | LFB | NA | 5681 | 2805 | 5293 | 932 | FAW13592 |
| 80.1 | DNA | R10 | GridIon | 457 | 1556 | NA | metagenomic | LFB | NA | 5468 | 2379 | 6421 | 1184 | FAX33641 |
| 81.1 | DNA | R10 | GridIon | 500 | 1733 | NA | metagenomic | LFB | NA | 7313 | 3103 | 5179 | 1068 | FAX57960 |
| 82.1 | DNA | R10 | GridIon | 507 | 1657 | NA | metagenomic | LFB | NA | 5308 | 2229 | 7428 | 1191 | FAX59252 |
| 83.1 | DNA | R10 | MinIon | 450 | 1657 | 156 | metagenomic | SFB | 600 | 7086 | 3361 | 1648 | 1199 | FAO79888 |
| 84.1 | DNA | R9 | MinIon | NA | 1953 | NA | metagenomic | SFB | 447 | NA | 7190 | 1355 | 2371 | FAK75664 |
| 85.1 | DNA | R9 | MinIon | NA | 1921 | NA | metagenomic | SFB | 471.6 | NA | 6634 | 1246 | 2878 | FAL78178 |
| 85.2 | DNA | R9 | MinIon | NA | 19 | NA | metagenomic | SFB | 471.6 | NA | NA | 836 | NA | FAL78178 |
| 86.1 | DNA | R9 | MinIon | NA | 186 | NA | metagenomic | SFB | 280 | NA | 555 | 2355 | NA | FAL78306 |
| 86.2 | DNA | R9 | MinIon | NA | 1915 | NA | metagenomic | SFB | 280 | NA | 7182 | 3067 | 3253 | FAL78306 |
| 87.1 | DNA | R9 | MinIon | 507 | 1933 | 154 | metagenomic | SFB | 252 | 17853 | 7006 | 2561 | 2339 | FAL80840 |
| 87.2 | DNA | R9 | MinIon | 396 | 1452 | 155 | metagenomic | SFB | 252 | 10310 | 5187 | 2378 | 1001 | FAL80840 |
| 88.1 | DNA | R9 | MinIon | 512 | NA | 167 | metagenomic | SFB | 256 | 19674 | 7000 | 2717 | 3098 | FAL81216 |
| 88.2 | DNA | R9 | MinIon | 378 | 1279 | 168 | metagenomic | SFB | 256 | 9050 | 4769 | 2434 | 1110 | FAL81216 |
| 89.1 | DNA | R9 | MinIon | 506 | 1793 | 147 | metagenomic | SFB | 301 | 6030 | 4453 | 1833 | 729 | FAL83332 |
| 89.2 | DNA | R9 | MinIon | 457 | 1617 | 148 | metagenomic | SFB | 301 | 2492 | 2204 | 1124 | 271 | FAL83332 |
| 90.1 | DNA | R9 | MinIon | 510 | 1966 | 179 | metagenomic | SFB | 1300 | 27326 | 6719 | 2119 | 2953 | FAL83351 |
| 91.1 | DNA | R9 | MinIon | 479 | 1813 | 177 | metagenomic | SFB | 431 | 13234 | 6063 | 1166 | 1671 | FAL84179 |
| 92.1 | DNA | R9 | MinIon | 485 | 1802 | 171 | metagenomic | SFB | 379 | 19643 | 5541 | 2215 | 2231 | FAL85078 |
| 93.1 | DNA | R9 | MinIon | 508 | NA | 150 | metagenomic | SFB | 360 | 14580 | 7302 | 4228 | 2041 | FAL91762 |
| 93.2 | DNA | R9 | MinIon | 420 | 1588 | 151 | metagenomic | SFB | 360 | 13564 | 5661 | 2288 | 1945 | FAL91762 |
| 94.1 | DNA | R9 | MinIon | 507 | 1891 | 181 | metagenomic | SFB | 400 | 17020 | 5792 | 1623 | 2302 | FAM93261 |
| 95.1 | DNA | R9 | MinIon | 376 | 1365 | 174 | metagenomic | SFB | 410 | 8949 | 4597 | 2509 | 1247 | FAM95301 |
| 96.1 | DNA | R9 | MinIon | 503 | 1843 | 190 | metagenomic | SFB | 609 | 25293 | 6351 | 1795 | 2686 | FAM95308 |
| 97.1 | DNA | R9 | MinIon | 508 | 1945 | 181 | metagenomic | SFB | 458 | 26777 | 7184 | 3221 | 2585 | FAM95383 |
| 98.1 | DNA | R9 | MinIon | 271 | 1005 | 174 | metagenomic | SFB | 300 | 2850 | 2272 | 2305 | 606 | FAM96150 |
| 99.1 | DNA | R9 | MinIon | 503 | 1896 | 178 | metagenomic | SFB | 564 | 17602 | 5838 | 1871 | 2520 | FAM96240 |
| 100.1 | DNA | R9 | MinIon | 465 | 1691 | 171 | metagenomic | SFB | 253 | 8206 | 5247 | 3313 | 1363 | FAM96327 |
| 100.2 | DNA | R9 | MinIon | 353 | 1293 | 172 | metagenomic | SFB | 253 | 6930 | 4180 | 1905 | 1291 | FAM96327 |
| 101.1 | DNA | R9 | MinIon | 507 | 1876 | 190 | metagenomic | SFB | 552 | 28264 | 6597 | 1462 | 3230 | FAM96767 |
| 102.1 | DNA | R9 | MinIon | 481 | 1780 | 54 | metagenomic | SFB | 460 | 12460 | 5585 | 1483 | 1076 | FAO10159 |
| 103.1 | DNA | R9 | MinIon | 499 | 1771 | 39 | metagenomic | SFB | 372 | 2181 | 1580 | 1713 | 812 | FAO10214 |
| 103.2 | DNA | R9 | MinIon | 240 | 796 | 45 | metagenomic | SFB | 460 | 3263 | 2633 | 1585 | 850 | FAO10214 |
| 104.1 | DNA | R9 | MinIon | 507 | 1594 | 40 | metagenomic | SFB | 960 | 4117 | 2129 | 922 | 1157 | FAO10253 |
| 105.1 | DNA | R9 | MinIon | 491 | 1602 | 41 | metagenomic | SFB | 408 | 2037 | 1597 | 1785 | NA | FAO10336 |
| 106.1 | DNA | R9 | MinIon | 498 | 1742 | 39 | metagenomic | SFB | 393 | 4078 | 2218 | 1444 | 1008 | FAO10413 |
| 10.5 | DNA | R9 | MinIon | 337 | 1157 | 55 | metagenomic | SFB | 564 | 3713 | 3571 | 1340 | 644 | FAO10418 |
| 107.1 | DNA | R9 | MinIon | 506 | 1948 | 42 | metagenomic | SFB | 634 | 23235 | 7055 | 1672 | 2661 | FAO12115 |
| 108.1 | DNA | R9 | MinIon | 510 | 1955 | 42 | metagenomic | SFB | 538 | 18522 | 6564 | 1740 | 2241 | FAO12140 |
| 109.1 | DNA | R9 | MinIon | 501 | 1881 | 42 | metagenomic | SFB | 600 | 18284 | 6171 | 1912 | 2292 | FAO12169 |
| 110.1 | DNA | R9 | MinIon | 484 | NA | 20 | metagenomic | SFB | 672 | 10957 | 5227 | 2273 | 1750 | FAO16442 |
| 111.1 | DNA | R9 | MinIon | 512 | 1853 | 48 | metagenomic | SFB | 240 | 19079 | 6678 | 1685 | 2428 | FAO83526 |
| 112.1 | DNA | R9 | MinIon | 509 | 1842 | 34 | metagenomic | SFB | 474 | 24501 | 6387 | 2150 | 3460 | FAO86805 |
| 113.1 | DNA | R9 | MinIon | 463 | 1637 | 87 | metagenomic | SFB | 189.6 | 3831 | 1777 | 2757 | 968 | FAP46095 |
| 114.1 | DNA | R9 | MinIon | 488 | 1659 | 87 | metagenomic | SFB | 158.4 | 15254 | 5286 | 1071 | 1976 | FAP49364 |
| 115.1 | DNA | R9 | MinIon | 500 | 1526 | 66 | metagenomic | SFB | 286.8 | 10070 | 3581 | 2934 | 1208 | FAQ01716 |
| 116.1 | DNA | R9 | MinIon | 479 | 1728 | 52 | metagenomic | SFB | 470.4 | 6142 | 2766 | 2240 | 834 | FAQ01897 |
| 117.1 | DNA | R9 | MinIon | 484 | 1680 | 52 | metagenomic | SFB | 477.6 | 7060 | 3049 | 1966 | 1301 | FAQ10186 |
| 118.1 | DNA | R9 | MinIon | 495 | 1850 | 36 | metagenomic | SFB | 674.4 | 12282 | 5140 | 1137 | 1761 | FAQ37458 |
| 119.1 | DNA | R9 | MinIon | 496 | 1817 | 37 | metagenomic | SFB | 574 | 11248 | 4596 | 1583 | 2062 | FAQ69372 |
| 120.1 | DNA | R9 | MinIon | 499 | 1904 | 151 | insect | LFB | 103 | 2509 | NA | 13240 | 264 | FAL78077 |
| 120.2 | DNA | R9 | MinIon | 481 | 1707 | 165 | insect | SFB | 113 | 5427 | 4328 | 5385 | 576 | FAL78077 |
| 121.1 | DNA | R9 | MinIon | 509 | 1984 | 150 | insect | LFB | 232 | 5127 | 4677 | 10627 | 438 | FAL83572 |
| 122.1 | DNA | R9 | MinIon | 380 | 1456 | 20 | insect | SFB | 255 | 2097 | 1977 | 5473 | 221 | FAO10283 |
| 10.1 | DNA | R9 | MinIon | 508 | NA | 49 | insect | LFB | 144 | 2393 | NA | 20800 | 386 | FAO10418 |
| 10.3 | DNA | R9 | MinIon | 390 | 1405 | 54 | insect | LFB | 144 | 1651 | NA | 21548 | 273 | FAO10418 |
| 10.4 | DNA | R9 | MinIon | 354 | 1257 | 55 | insect | SFB | 255 | 1084 | NA | 5500 | 162 | FAO10418 |
| 123.1 | DNA | R9 | MinIon | 462 | 1666 | 7 | insect | SFB | 226 | 7190 | 5361 | 4274 | 729 | FAO10430 |
| 123.2 | DNA | R9 | MinIon | 373 | 1394 | 21 | insect | SFB | 255 | 1473 | 1441 | 10126 | 165 | FAO10430 |
| 124.1 | DNA | R9 | MinIon | 497 | 1843 | 69 | insect | LFB | 200 | 2766 | 2638 | 16530 | 263 | FAO12153 |
| 124.2 | DNA | R9 | MinIon | 447 | 1699 | 74 | insect | LFB | 200 | 2253 | 2205 | 16834 | 244 | FAO12153 |
| 124.3 | DNA | R9 | MinIon | 388 | 1512 | 75 | insect | LFB | 160 | 1771 | NA | 15219 | 229 | FAO12153 |
| 124.4 | DNA | R9 | MinIon | 358 | 1433 | 76 | insect | LFB | 160 | 1413 | NA | 16750 | 226 | FAO12153 |
| 124.5 | DNA | R9 | MinIon | 355 | 1340 | 88 | insect | LFB | 150 | 1716 | NA | 19073 | 249 | FAO12153 |
| 124.6 | DNA | R9 | MinIon | 230 | 951 | 89 | insect | LFB | 100 | 1003 | NA | 14562 | 258 | FAO12153 |
| 125.1 | DNA | R9 | MinIon | 436 | 1540 | 60 | insect | LFB | 300 | 3879 | 3655 | 16927 | 361 | FAO12159 |
| 125.2 | DNA | R9 | MinIon | 410 | 1489 | 61 | insect | LFB | 200 | 2721 | 2671 | 18638 | 272 | FAO12159 |
| 125.3 | DNA | R9 | MinIon | 416 | 1591 | 61 | insect | LFB | 200 | 3097 | NA | 13451 | 547 | FAO12159 |
| 125.4 | DNA | R9 | MinIon | 364 | 1366 | 62 | insect | LFB | 200 | 2271 | NA | 14192 | 338 | FAO12159 |
| 125.5 | DNA | R9 | MinIon | 321 | 1196 | 67 | insect | LFB | 200 | 1319 | NA | 16557 | 175 | FAO12159 |
| 125.6 | DNA | R9 | MinIon | 264 | 1086 | 68 | insect | LFB | 100 | 719 | NA | 16788 | 151 | FAO12159 |
| 126.1 | DNA | R9 | MinIon | 510 | 1935 | 49 | insect | SFB | 200 | 9855 | 6485 | 4358 | 1016 | FAO86712 |
| 126.2 | DNA | R9 | MinIon | 476 | 1811 | 50 | insect | SFB | 100 | 6855 | 5140 | 4640 | 724 | FAO86712 |
| 126.3 | DNA | R9 | MinIon | 419 | 1570 | 51 | insect | SFB | 67 | 4758 | 4070 | 4859 | 545 | FAO86712 |
| 126.4 | DNA | R9 | MinIon | 307 | 1381 | 62 | insect | LFB | 70 | 1701 | 1587 | 7019 | 264 | FAO86712 |
| 127.1 | DNA | R9 | MinIon | 445 | 1709 | 8 | insect | LFB | 200 | 4467 | 4156 | 18700 | 443 | FAO86834 |
| 127.2 | DNA | R9 | MinIon | 468 | 1785 | 9 | insect | LFB | 170 | 3610 | 3305 | 18665 | 336 | FAO86834 |
| 128.1 | DNA | R9 | MinIon | 383 | 1424 | 6 | insect | LFB | 200 | 1543 | 1510 | 19752 | 113 | FAO88446 |
| 129.1 | DNA | R9 | MinIon | 487 | 1796 | 55 | insect | LFB | 190 | 4379 | 3694 | 6359 | 357 | FAO88464 |
| 129.2 | DNA | R9 | MinIon | 279 | 1046 | 57 | insect | LFB | 50 | 1507 | NA | 6788 | 311 | FAO88464 |
| 130.1 | DNA | R9 | MinIon | 511 | 1944 | 44 | insect | SFB | 200 | 9475 | 6638 | 4590 | 922 | FAO91712 |
| 130.2 | DNA | R9 | MinIon | 464 | 1753 | 48 | insect | SFB | 100 | 6429 | 4884 | 4961 | 692 | FAO91712 |
| 130.3 | DNA | R9 | MinIon | 425 | NA | 51 | insect | SFB | 67 | 2592 | 2592 | 4828 | 525 | FAO91712 |
| 67.1 | DNA | R9 | MinIon | 499 | 1907 | 8 | insect | LFB | 200 | 4891 | 4586 | 18872 | 455 | FAO92001 |
| 67.2 | DNA | R9 | MinIon | 485 | 1852 | 9 | insect | LFB | 170 | 4284 | 3895 | 19414 | 417 | FAO92001 |
| 67.3 | DNA | R9 | MinIon | 473 | 1776 | 13 | insect | LFB | 140 | 3997 | 3559 | 18674 | 368 | FAO92001 |
| 67.5 | DNA | R9 | MinIon | 427 | 1508 | 44 | insect | SFB | 78 | 4948 | 3728 | 3529 | 607 | FAO92001 |
| 67.6 | DNA | R9 | MinIon | 111 | 758 | 51 | insect | LFB | 30 | 648 | 578 | 14885 | 408 | FAO92001 |
| 131.1 | DNA | R9 | MinIon | 512 | 1959 | 55 | insect | LFB | 190 | 7742 | 6090 | 6314 | 803 | FAO92100 |
| 131.2 | DNA | R9 | MinIon | 498 | 1864 | 57 | insect | LFB | 100 | 3275 | NA | 6876 | NA | FAO92100 |
| 131.3 | DNA | R9 | MinIon | 469 | 1772 | 57 | insect | LFB | 85 | 5929 | 4737 | 4932 | 894 | FAO92100 |
| 131.4 | DNA | R9 | MinIon | 380 | 1421 | 58 | insect | LFB | 60 | 3028 | 2746 | 6700 | 370 | FAO92100 |
| 131.5 | DNA | R9 | MinIon | 223 | 874 | 62 | insect | LFB | 70 | 807 | 778 | 6584 | 264 | FAO92100 |
| 132.1 | DNA | R9 | MinIon | 465 | 1812 | 57 | insect | LFB | 100 | 4724 | 4089 | 7116 | 498 | FAO92220 |
| 132.2 | DNA | R9 | MinIon | 359 | 1379 | 58 | insect | LFB | 60 | 2643 | 2390 | 6769 | 334 | FAO92220 |
| 133.1 | DNA | R9 | MinIon | 506 | 1951 | 58 | insect | LFB | 190 | 6564 | 5533 | 6795 | 688 | FAO92291 |
| 133.2 | DNA | R9 | MinIon | 433 | 1628 | 62 | insect | LFB | 85 | 3866 | 3453 | 7461 | 426 | FAO92291 |
| 134.1 | DNA | R9 | MinIon | 488 | 1725 | 72 | plant | SFB | 350 | 8694 | 3799 | 2288 | 620 | FAR96897 |
| 135.1 | DNA | R9 | MinIon | 473 | 1615 | 72 | plant | SFB | 330 | 9350 | 3929 | 4101 | 532 | FAR96905 |
| 136.1 | RNA | R9 | MinIon | 511 | 1399 | 128 | mouse | RNA | 1252 | 1407 | 910 | 1058 | 915 | FAO83512 |
| 137.1 | RNA | R9 | MinIon | 509 | 1383 | 128 | mouse | RNA | 1372 | 832 | 705 | 746 | 638 | FAO92305 |
| 138.1 | RNA | R9 | MinIon | 494 | 1275 | 12 | mouse | RNA | 1092 | 1206 | 680 | 733 | 1094 | FAP46096 |
| 139.1 | RNA | R9 | MinIon | 474 | 1263 | 12 | mouse | RNA | 1252 | 1174 | 708 | 772 | 1176 | FAP46400 |
| 140.1 | RNA | R9 | MinIon | 501 | 1353 | 12 | mouse | RNA | 1220 | 1940 | 888 | 1049 | 1390 | FAP46452 |
| 141.1 | RNA | R9 | MinIon | 486 | 1320 | 12 | mouse | RNA | 1044 | 1337 | 800 | 1091 | 827 | FAP49780 |
| 142.1 | DNA | R9 | MinIon | 494 | NA | 86 | mouse | SFB | 385 | 10671 | 6305 | 10446 | 1935 | FAQ58673 |
| 142.2 | DNA | R9 | MinIon | NA | 1236 | NA | mouse | SFB | 385 | NA | 2280 | 9714 | 370 | FAQ58673 |
| 143.1 | DNA | R9 | MinIon | 479 | 1827 | 63 | mouse | SFB | 300 | 6680 | 4769 | 8821 | 599 | FAQ58674 |
| 144.1 | DNA | R9 | MinIon | 447 | 1714 | 57 | mouse | SFB | 300 | 5561 | 4246 | 9428 | 612 | FAQ61173 |
| 145.1 | DNA | R9 | MinIon | 499 | NA | 83 | mouse | SFB | 245 | 11137 | 6360 | 10034 | 1800 | FAQ61176 |
| 145.2 | DNA | R9 | MinIon | NA | 1205 | NA | mouse | SFB | 245 | NA | 2779 | 9350 | 510 | FAQ61176 |
| 146.1 | DNA | R9 | MinIon | 508 | 1949 | 63 | mouse | SFB | 282 | 21067 | 6967 | 9473 | 1993 | FAQ69208 |
| 147.1 | DNA | R9 | MinIon | 508 | 1875 | 83 | mouse | SFB | 245 | 19740 | 6786 | 9155 | 1855 | FAQ69274 |
| 148.1 | DNA | R9 | MinIon | 499 | 1909 | 83 | mouse | SFB | 385 | 19034 | 6723 | 9377 | 1832 | FAQ69338 |
| 149.1 | DNA | R9 | MinIon | 507 | 1895 | 49 | mouse | SFB | 270 | 24681 | 7030 | 6962 | 2518 | FAQ69370 |
| 150.1 | DNA | R9 | MinIon | 490 | 1829 | 57 | mouse | SFB | 280 | 22628 | 6920 | 8963 | 2237 | FAQ69578 |
| 151.1 | DNA | R9 | MinIon | 509 | 1964 | 170 | human | SFB | 444 | 18354 | 6760 | 1550 | 2234 | FAL84151 |
| 151.2 | DNA | R9 | MinIon | 213 | 728 | 172 | human | SFB | 444 | 2207 | 1506 | 1485 | 737 | FAL84151 |
| 152.1 | RNA | R9 | MinIon | 505 | 1360 | 47 | human | RNA | 42 | 157 | 117 | 475 | 431 | FAO10420 |
| 153.1 | DNA | R9 | MinIon | 502 | NA | 68 | human | SFB | 538 | 12217 | 6859 | 2414 | 1762 | FAP46521 |
| 154.1 | DNA | R9 | MinIon | 489 | NA | 43 | human | SFB | 451 | 10568 | 6294 | 4487 | 1612 | FAP49434 |
| 154.2 | DNA | R9 | MinIon | 448 | 1708 | 44 | human | SFB | 451 | 13249 | 5635 | 4181 | 1369 | FAP49434 |
| 155.1 | DNA | R9 | MinIon | NA | 1919 | NA | human | SFB | 688.8 | NA | 6763 | 4626 | 2319 | FAP49725 |
| 156.1 | DNA | R9 | MinIon | 501 | 1928 | 189 | human | LFB | 400 | 12833 | NA | 5179 | 1386 | FAQ01752 |
| 157.1 | DNA | R9 | MinIon | 467 | 1430 | 57 | human | SFB | 382 | 7352 | 4147 | 515 | 712 | FAQ09810 |
| 158.1 | DNA | R9 | MinIon | 480 | 1356 | 49 | human | SFB | 821 | 6529 | 4541 | 454 | 899 | FAQ10598 |
| 158.2 | DNA | R9 | MinIon | 187 | 536 | 50 | human | SFB | 547 | 2326 | 1799 | 467 | 791 | FAQ10598 |
| 158.3 | DNA | R9 | MinIon | 76 | 239 | 51 | human | SFB | 274 | 299 | 227 | 854 | NA | FAQ10598 |
| 159.1 | DNA | R9 | MinIon | 488 | 1878 | 162 | human | SFB | 300 | 13259 | 5853 | 2776 | 1275 | FAQ69371 |
| 160.1 | DNA | R9 | MinIon | 476 | NA | 92 | human | SFB | 400 | 15275 | 5655 | 2746 | 1314 | FAR29925 |
| 161.1 | DNA | R9 | MinIon | NA | 1690 | 238 | human | LFB | 405 | NA | 4401 | 2257 | 544 | FAR33296 |
| 161.2 | DNA | R9 | MinIon | 293 | 866 | NA | human | LFB | 405 | 2509 | 1905 | 927 | 350 | FAR33296 |
| 162.1 | DNA | R9 | MinIon | 509 | NA | 123 | human | LFB | 394 | 19595 | NA | NA | 1491 | FAR33573 |
| 163.1 | DNA | R9 | MinIon | 456 | NA | 130 | human | LFB | 305 | 8441 | NA | 643 | 759 | FAR38117 |
| 164.1 | DNA | R9 | MinIon | 505 | NA | 27 | human | LFB | 365 | 9027 | NA | NA | 935 | FAR39219 |
| 165.1 | DNA | R9 | MinIon | 498 | NA | 37 | human | LFB | 321 | 7285 | NA | 765 | 750 | FAR39290 |
| 166.1 | DNA | R9 | MinIon | 508 | NA | 74 | human | LFB | 453 | 6073 | NA | 944 | 834 | FAR39317 |
| 167.1 | DNA | R9 | MinIon | 377 | 1197 | 242 | human | LFB | 340 | 3463 | 2744 | 677 | 356 | FAR39502 |
| 168.1 | DNA | R9 | MinIon | 490 | NA | 91 | human | SFB | 400 | 17476 | 5708 | 4673 | 1212 | FAR93058 |
| 169.1 | DNA | R9 | MinIon | 383 | 993 | 247 | human | LFB | 340 | 2266 | 1726 | 848. | 221 | FAR96341 |
| 170.1 | DNA | R9 | MinIon | 481 | 1833 | 92 | human | SFB | 400 | 15910 | 5763 | 5055 | 1078 | FAR96408 |
| 171.1 | DNA | R9 | MinIon | 483 | 1777 | 91 | human | SFB | 400 | 10105 | 4151 | 2782 | 1001 | FAR96434 |
| 172.1 | DNA | R9 | MinIon | 384 | 1080 | 249 | human | LFB | 425 | 3106 | 2509 | 1029 | 397 | FAR96514 |
| 173.1 | DNA | R9 | MinIon | 510 | 1882 | 37 | human | SFB | 300 | 7049 | 5165 | 1412 | 718 | FAR96669 |
| 173.2 | DNA | R9 | MinIon | 300 | 1079 | 38 | human | SFB | 200 | 5071 | 2886 | 1414 | 726 | FAR96669 |
| 174.1 | DNA | R9 | MinIon | 479 | 1294 | 238 | human | LFB | 389 | 5313 | 3079 | 849 | 446 | FAR96893 |
| 175.1 | DNA | R9 | MinIon | 438 | 1308 | 171 | human | LFB | 405 | 3491 | 2671 | 947 | 630 | FAS60674 |
| 176.1 | DNA | R9 | MinIon | 509 | NA | 90 | human | LFB | 450 | 20857 | 6391 | 587 | 2090 | FAV38963 |
| 176.2 | DNA | R9 | MinIon | NA | 728 | 91 | human | LFB | 411 | NA | 1729 | 1164 | 584 | FAV38963 |
| 177.1 | DNA | R9 | MinIon | NA | NA | 69 | human | LFB | 400 | NA | NA | NA | NA | FAV38989 |
| 177.2 | DNA | R9 | MinIon | 452 | 1445 | NA | human | LFB | 330 | 8250 | 4531 | 907 | 984 | FAV38989 |
| 178.1 | DNA | R9 | MinIon | 511 | NA | 60 | human | LFB | 400 | 14634 | 5579 | 622 | 1222 | FAV39381 |
| 179.1 | DNA | R9 | MinIon | NA | 1437 | NA | human | LFB | 412 | NA | 6287 | 679 | 1644 | FAV87040 |
| 180.1 | DNA | R9 | MinIon | NA | 1371 | NA | human | LFB | 414.96 | NA | 5864 | 569 | 1207 | FAW80737 |
| 9.2 | RNA | R9 | MinIon | 270 | 694 | 83 | synthetic | RNA | 230 | 505 | 362 | 893 | 801 | FAO07649 |
| 124.7 | RNA | R9 | MinIon | 189 | 484 | 103 | synthetic | RNA | 207 | 201 | 183 | 1002 | 469 | FAO12153 |
| 125.7 | RNA | R9 | MinIon | 200 | 522 | 103 | synthetic | RNA | 212 | 195 | 179 | 1038 | 418 | FAO12159 |

**Figure S1.**
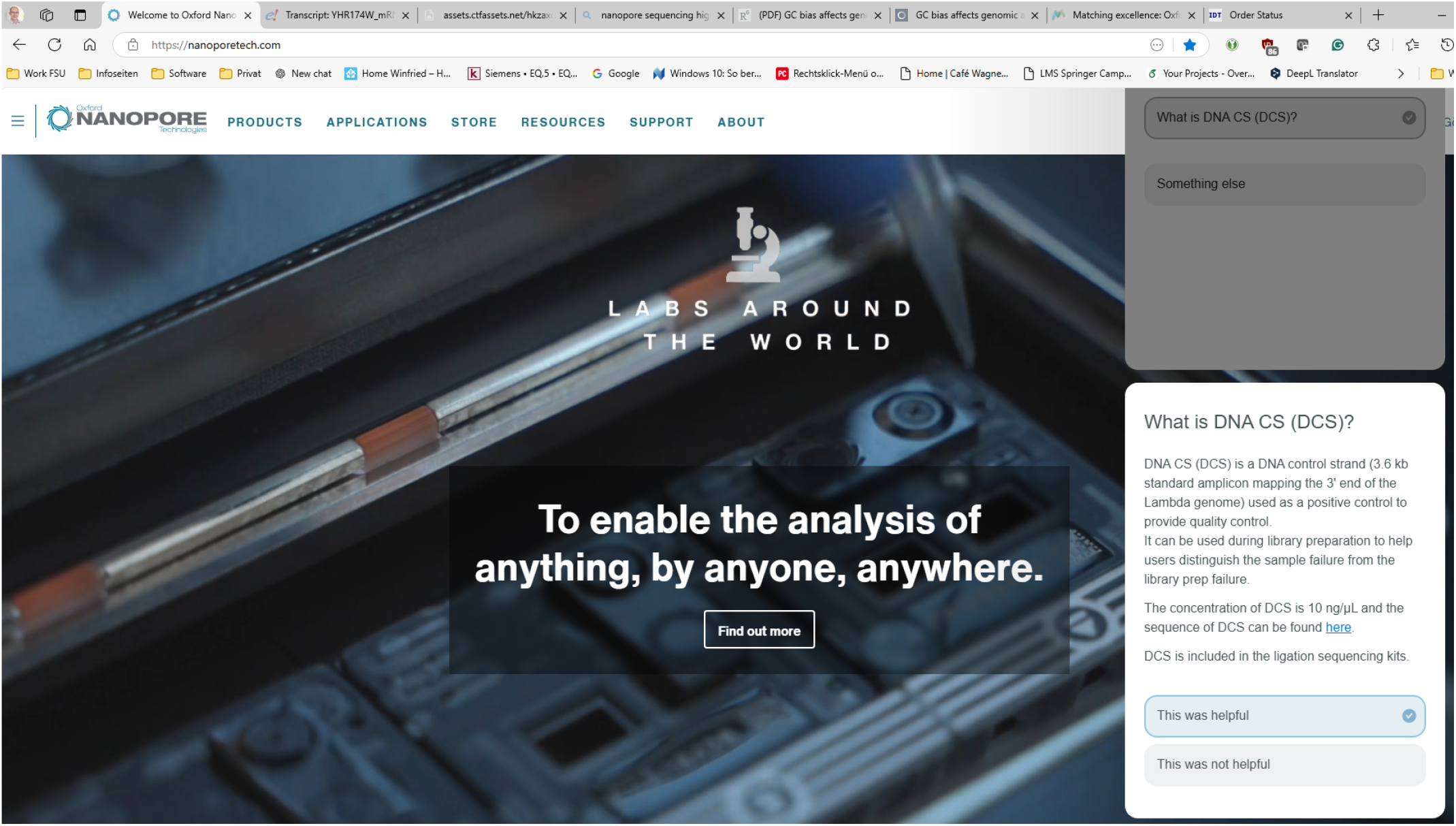
Screenshot from the Nanopore Community about DNA control sequence (DCS)

### ACCESS RAW ONT DATA

The following equation is used to convert the squiggles stored in the raw data to the pA signal:

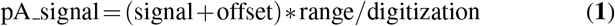

where `signal` represents the raw integer value, `offset` is the shift in the pA signal, `range` is the range of the pA signal, and `digitization` is the resolution of the conversion used to convert the continuous pA signal into discrete integer values. All parameters are provided by ONT in the raw data files.

### NORMALIZATION

Normalization corrects for biases, ensuring that signals from different experiments or sequencing channels are directly comparable. It can be performed on each read individually using statistical measures such as the mean and standard deviation or the median and median absolute deviation (MAD). The median and MAD are more robust to outliers, which are common in sequencing data, see Fig. 10. ONT tools, including their basecallers, typically employ normalization during data processing. The standard formula for signal normalization, based on the median and MAD, is as follows:

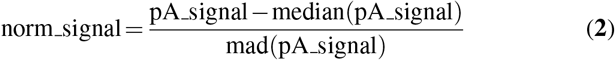

This process ensures that the signal distributions across reads become uniform, eliminating biases (see Fig. **??**B).

**Table S2.** Overview of methylation calling tools (adapted from Liu *et al*.), sorted by publication date. 5mC – 5-methylcytosine in DNA; m5C – 5-methylcytosine in RNA; 6mA – N6-methyladenosine in DNA; m6A – N6-methyladenosine in RNA; 5hmC – 5-hydroxymethylcytosine; psU – pseudouridine; m1A – N1-methyladenosine; 5moU – 5-methoxyuridine; m7G – N7-methylguanosine; Ino – inosine; f5C – 5-formylcytidine;

| comp. approach | tool | DNA / RNA | modification type | year | citation |
| --- | --- | --- | --- | --- | --- |
| hidden Markov models | Nanopolish | DNA | 5mC | 2017 | (79) |
| statistical | Tombo | DNA / RNA | 4mC, 5mC, m5C, 6mA | 2017 | (86) |
| statistical | SignalAlign | DNA | 5mC, 5hmC, 6mA | 2017 | (87) |
| neural networks | Dorado | DNA | 5mC, 6mA |  | ONT<sup>1</sup> |
| statistical | NanoMod | DNA | 5mC | 2019 | (88) |
| neural networks | mCaller | DNA | 6mA | 2019 | (89) |
| neural networks | DeepSignal | DNA | 5mC, 6mA | 2019 | (90) |
| neural networks | DeepMod | DNA | 5mC, 6mA | 2019 | (91) |
| neural network | DeepMod2 | DNA | 5mC | 2024 | (92) |
| neural networks | Megalodon | DNA | 5mC, 6mA |  | ONT<sup>2</sup> |
| hidden Markov models | f5c | DNA | 5mC | 2020 | (75) |
| neural networks | methBERT | DNA | 5mC, 6mA | 2021 | (93) |
| meta approaches | METEORE | DNA | 5mC, 6mA | 2021 | (21) |
| neural networks | DeepMP | DNA | 5mC, 6mA | 2021 | (94) |
| statistical | – | DNA | D2O | 2024 | (85) |
| machine learning | MINES | RNA | m6A | 2019 | (95) |
| error profile & machine learning | EpiNano | RNA | m6A, psU | 2019 | (96) |
| neural networks | nanoDoc | RNA | m6A | 2020 | (97) |
| neural networks | nano-ID | RNA | e5U | 2020 | (98) |
| statistical | DRUMMER | RNA | m6A | 2020 | (99) |
| statistical | xPore | RNA | m6A | 2021 | (100) |
| machine learning | Nanom6A | RNA | m6A | 2021 | (101) |
| statistical | Yanocomp | RNA | m6A | 2021 | (?) |
| machine learning | nanoRMS | RNA | psU | 2021 | (102) |
| error profile | ELIGOS | RNA | m6A, m1A, 5moU, psU, m7G, Ino, hm5C, f5C | 2021 | (103) |
| statistical | Nanocompore | RNA | m6A | 2021 | (104) |
| neural networks | m6Anet | RNA | m6A | 2022 | (105) |
| machine learning | Penguin | RNA | psU | 2022 | (106) |
| neural networks | DENA | RNA | m6A | 2022 | (107) |
| statistical | Magnipore | RNA | - | 2023 | (108) |
| neural networks | mAFIA | RNA | m6A | 2024 | (109) |

<sup>1</sup> https://github.com/nanoporetech/dorado
<sup>2</sup> https://github.com/nanoporetech/megalodon

**Figure S2.**
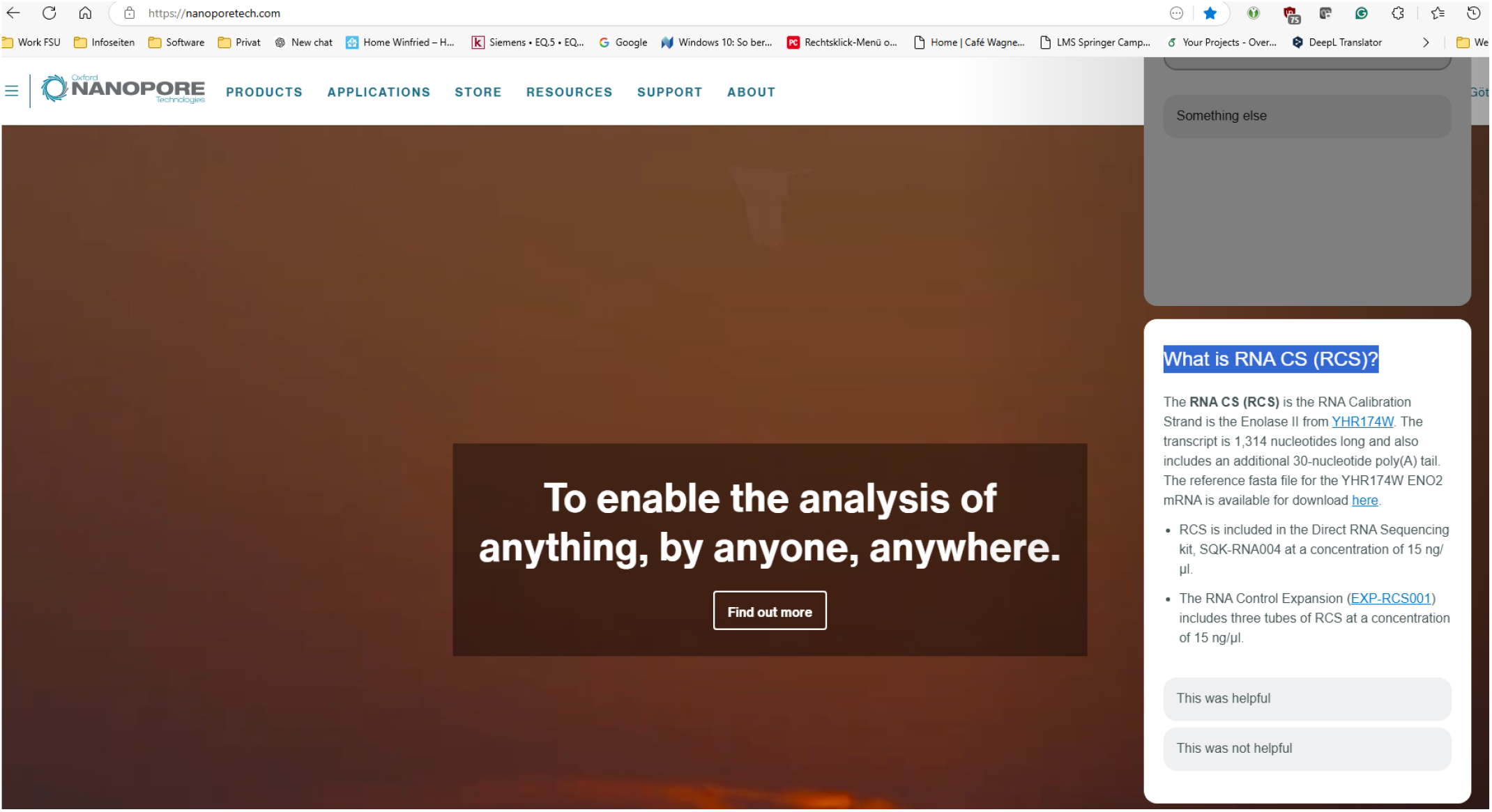
Screenshot from the Nanopore Community about RNA control sequence (RCS)

**Figure S3.**
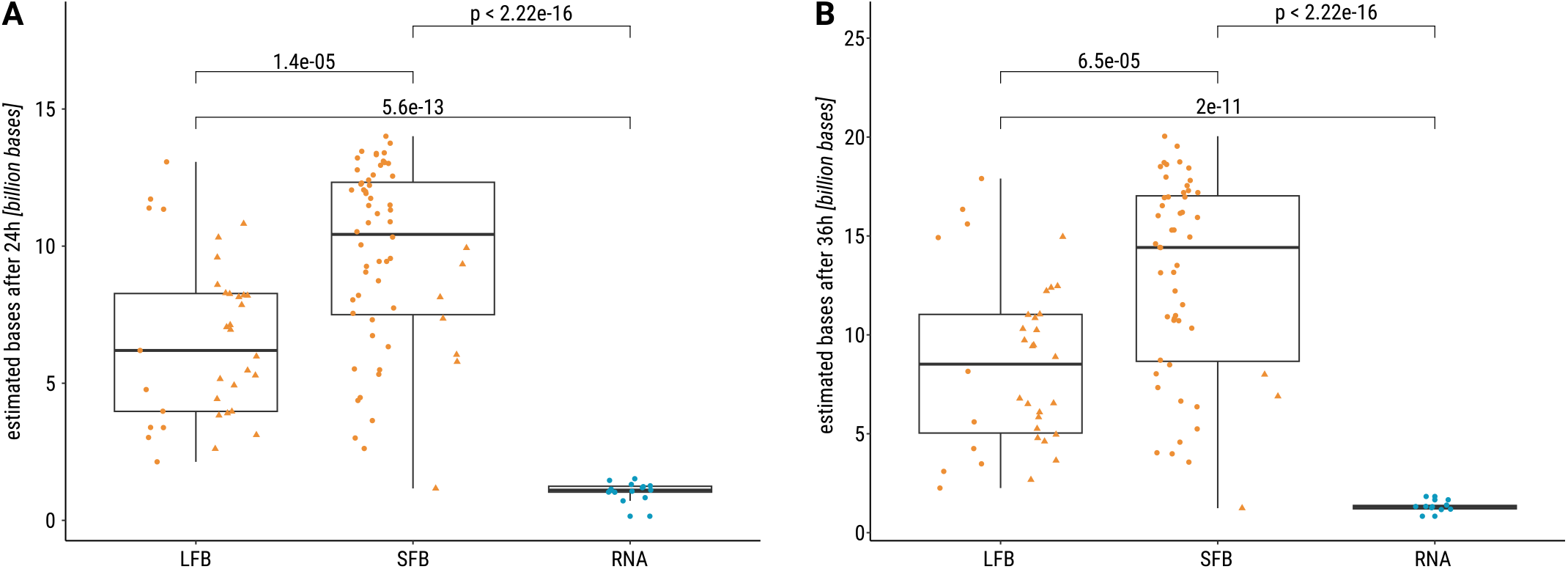
T-test shows higher estimated number of sequenced bases using SFB compared to LFB buffer after 24 h (**A**) and 36 h (**B**); addition to Fig. 4.

**Figure S4.**
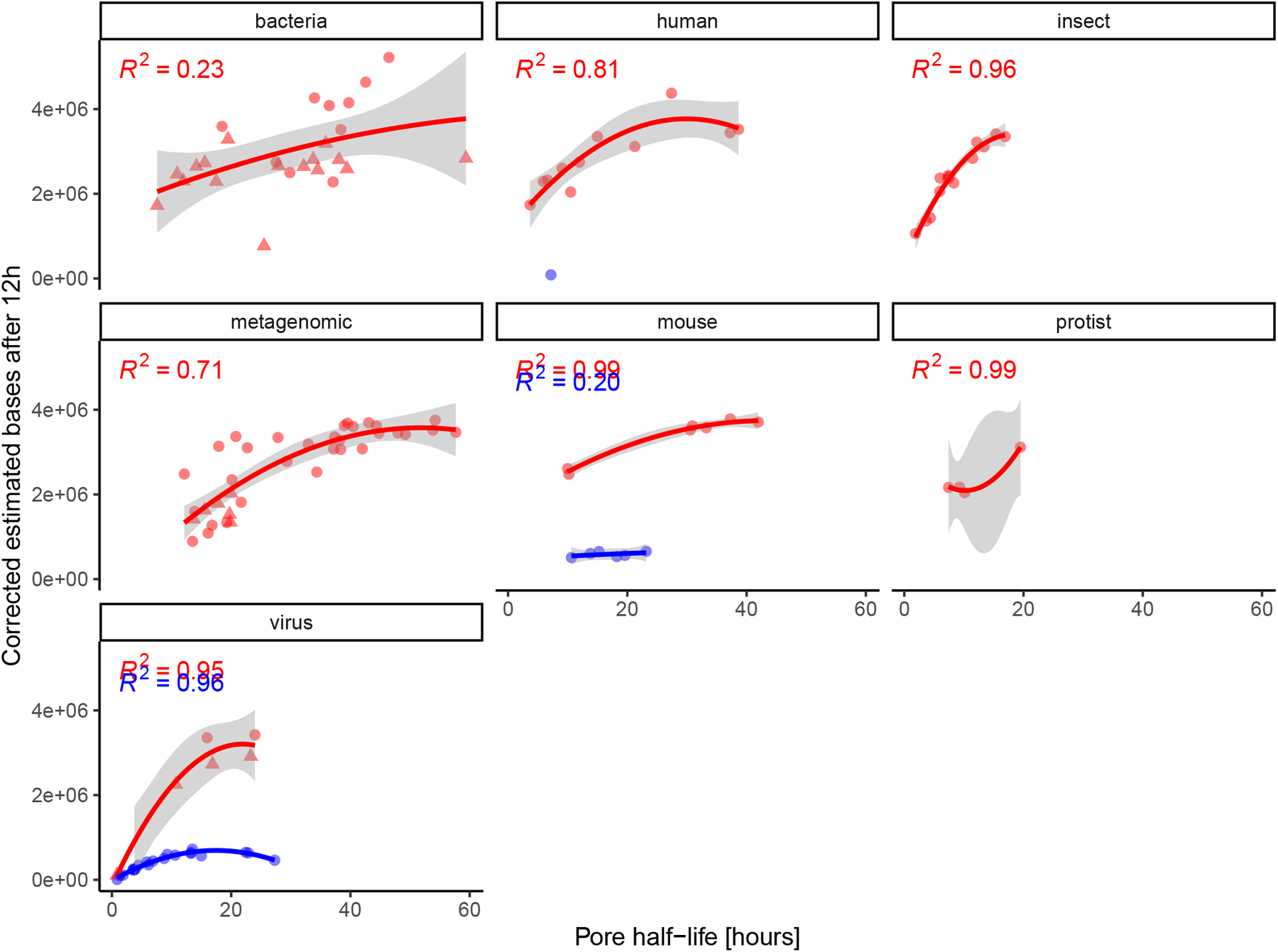
Correlation (R^2^) between pore half life and the estimated number of bases after 12 hours of sequencing (EB12) is observed across most sample types. The EB12 values were divided by number of starting pores, to corrected for any potential bias. The resulting plots reflect the same statements as Fig. 6A.

**Figure S5.**
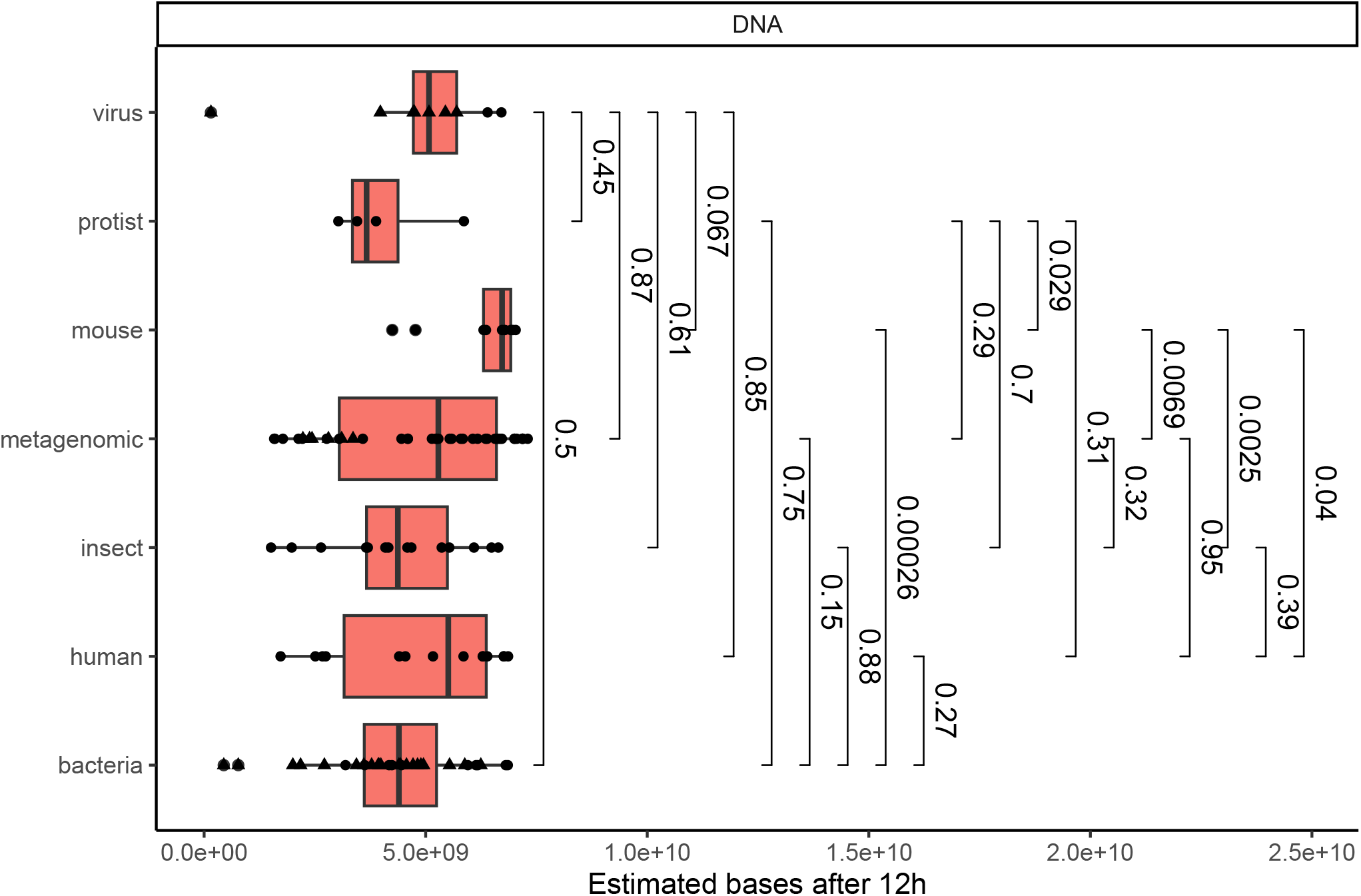
T-test shows higher EB12 in mouse compared to other sample types, addition to Fig. 6B.

**Figure S6.**
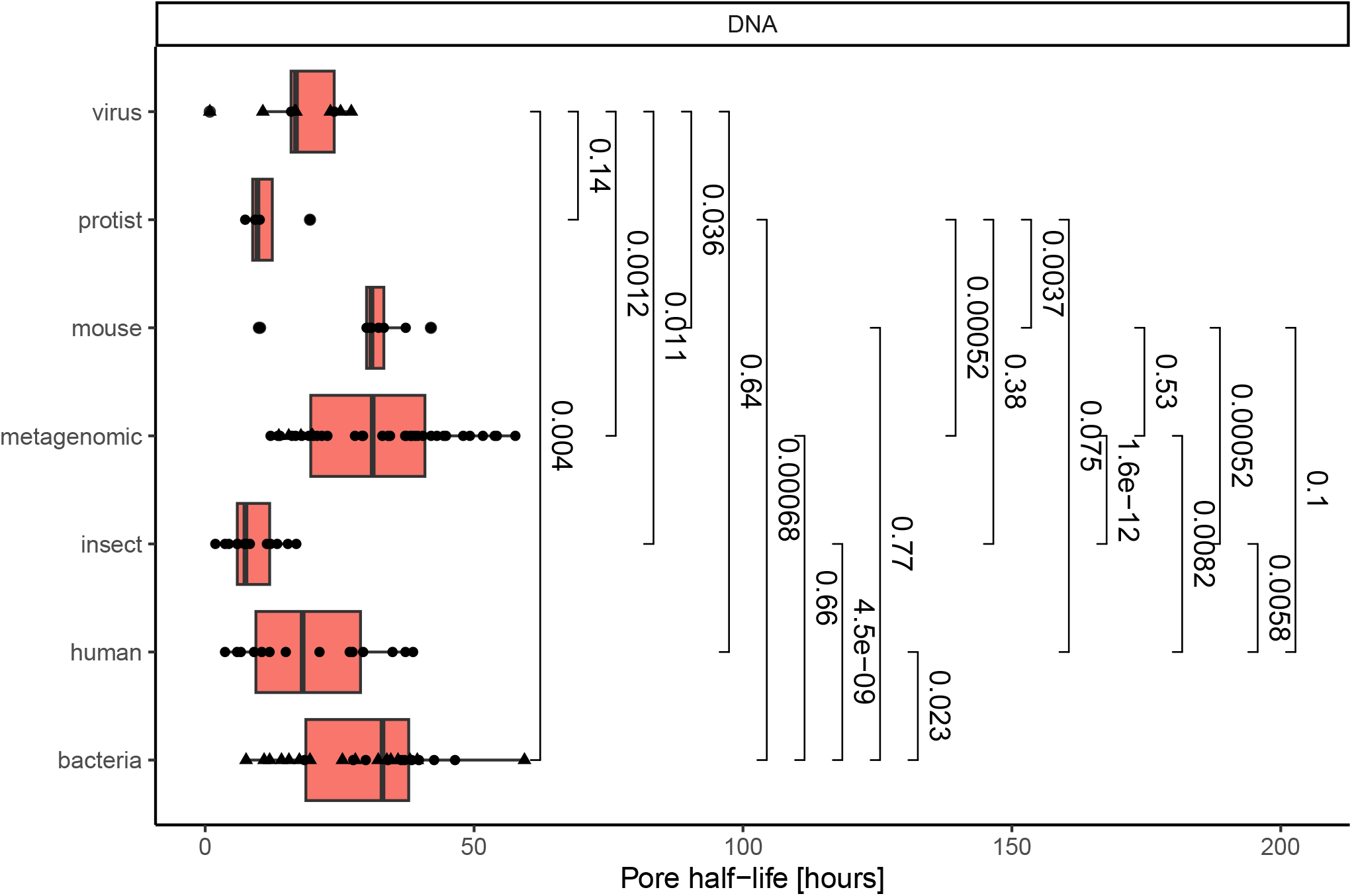
T-test shows significant differences in pore half time between sample types; addition to Fig. 6C.

**Figure S7.**
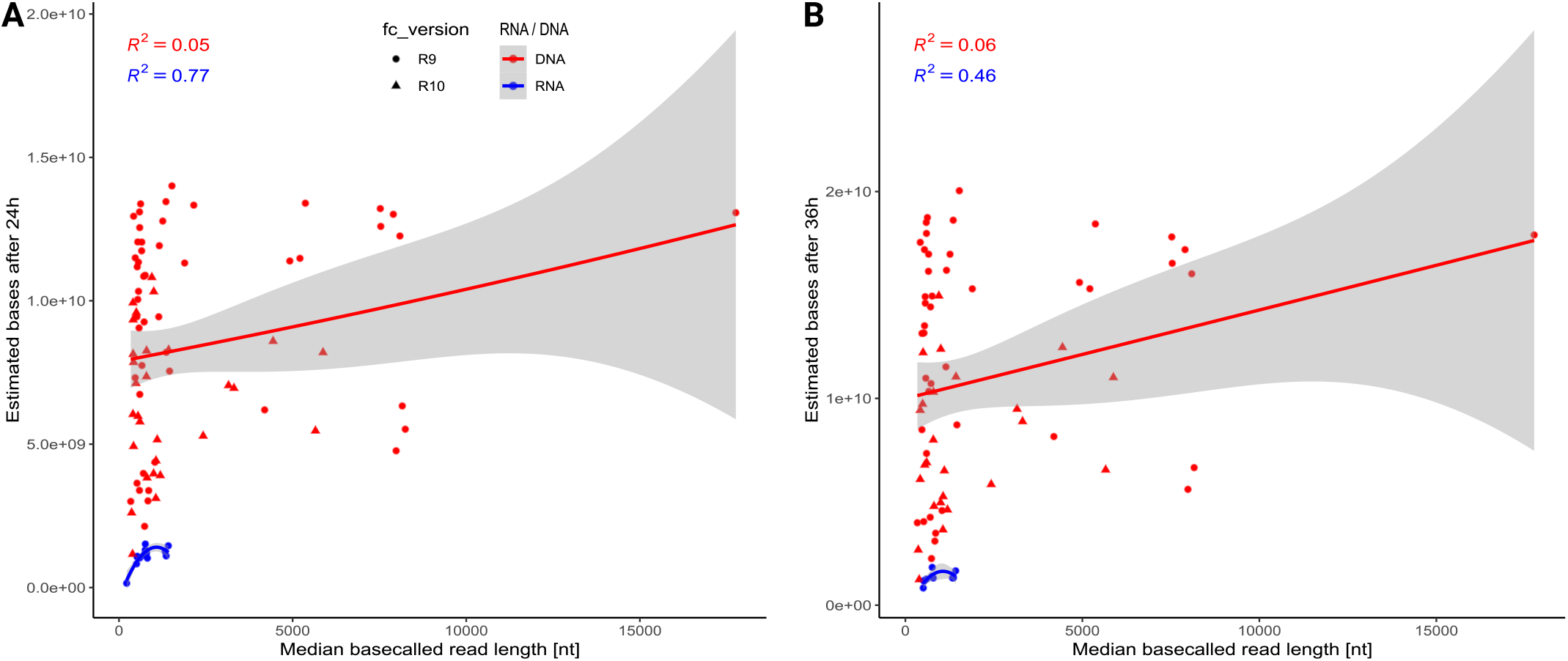
Distribution of estimated number of sequenced bases after 24 h (**A**) and 36 h (**B**) for DNA (red) and RNA (blue) samples; addition to Fig. 6D.

**Table S3.** Re-evaluated hand picked pore half time for Different Run IDs.

| ID | Half Life |
| --- | --- |
| 1.1 | 998 |
| 2.1 | 1630 |
| 6.1 | 1510 |
| 10.1 | 386 |
| 88.1 | 3098 |
| 93.1 | 2041 |
| 110.1 | 1750 |
| 130.3 | 525 |
| 142.1 | 1935 |
| 145.1 | 1800 |
| 153.1 | 1762 |
| 154.1 | 1612 |
| 160.1 | 1314 |
| 162.1 | 1491 |
| 163.1 | 759 |
| 164.1 | 935 |
| 165.1 | 750 |
| 166.1 | 834 |
| 168.1 | 1212 |
| 178.1 | 1222 |
| 176.1 | 1971 |
| 176.1 | 2090 |

**Figure S8.**
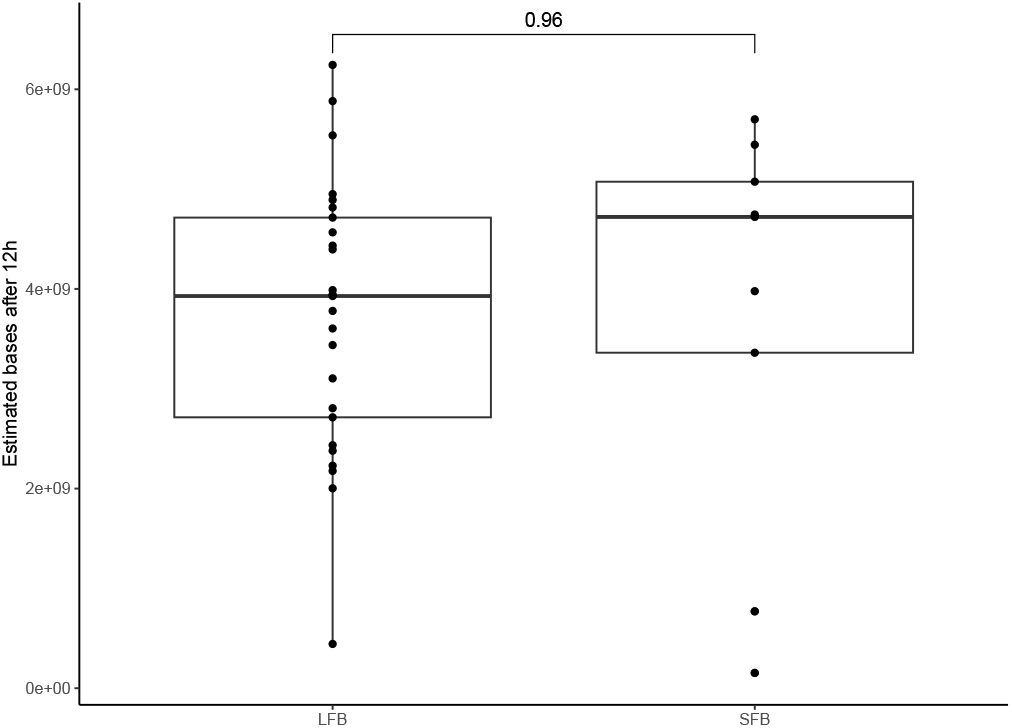
T-test shows no significant differences between the buffers used for R10 data. This might be due to the low number of data points.

**Table S4.**
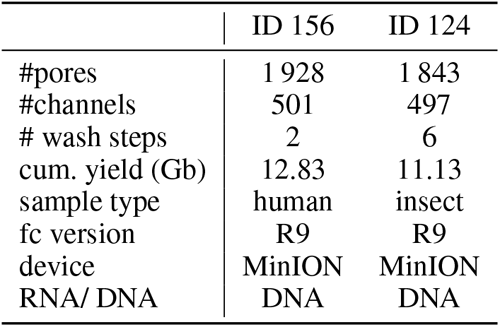
Sequencing statistics for the samples used in Fig. 7. cum. yield (Gb) – cumulative amount of estimated bases for all sequencing runs on this flow cell. done

**Figure S9.**
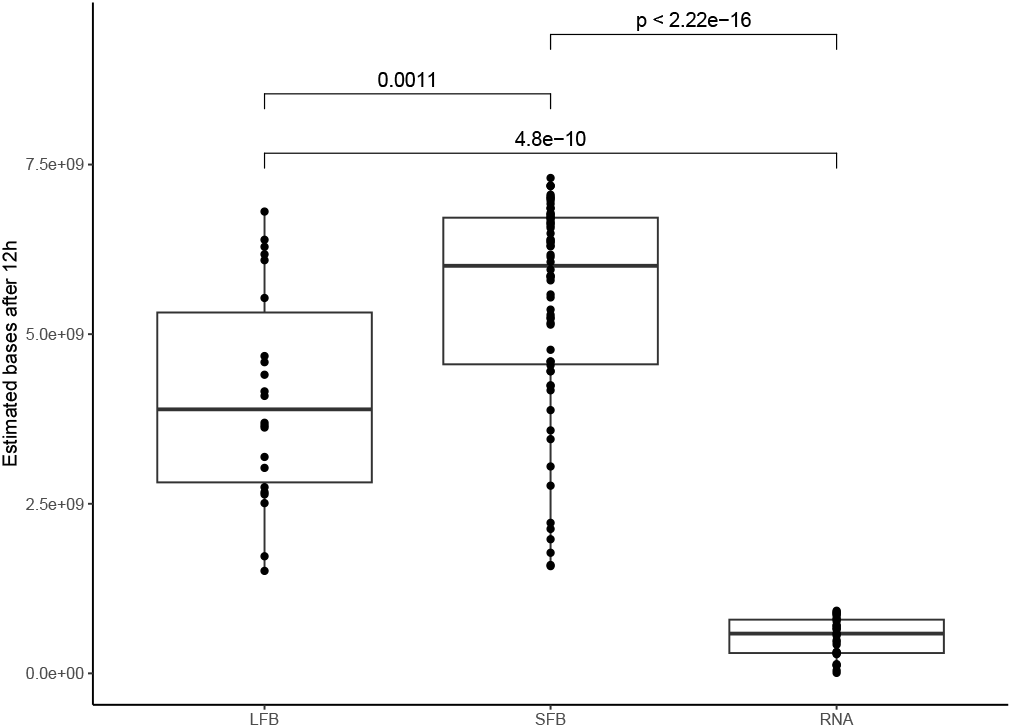
T-test shows significant differences between the buffers used for R9 data.

**Table S5.** Samples used for adaptive sampling in Fig. 8. # por – amount of active pores at sequencing start; yield – cumulative sequencing yield in Gb; depth – mean sequencing depth on CpG islands; wash – number of times the flow cell was washed in between; buffer – extension length of the CpG islands

| id | # por | yield | depth | wash | buffer |
| --- | --- | --- | --- | --- | --- |
| NA | NA | 10.7 | 3.01 | 0 | 0 |
| NA | 1519 | 19.7 | 6.10 | 0 | 0 |
| 155 | 1504 | 16.7 | 4.75 | 0 | 0 |
| 154 | 1305 | 23.3 | 6.67 | 1 | 0 |
| 153 | 1379 | 11.4 | 3.28 | 0 | 1 |
| 153 | 822 | 2.58 | 1.39 | 0 | 1 |
| NA | 1001 | 2.59 | 0.67 | 0 | 500 |
| 158 | 1192 | 8.65 | 2.13 | 2 | 500 |
| 157 | 1269 | 8.72 | 4.76 | 1 | 500 |
| 156 | 1250 | 13.49 | 3.85 | 2 | 0 |
| 164 | 1405 | 9.68 | 8.79 | 1 | 2000 |
| 165 | 1433 | 7.74 | 8.88 | 2 | 2000 |
| 166 | 1548 | 6.39 | 7.60 | 2 | 2000 |
| 162 | 1620 | 21.26 | 17.79 | 1 | 2000 |
| 163 | 969 | 9.00 | 7.24 | 1 | 2000 |
| 174 | 1352 | 6.46 | 5.40 | 1 | 2000 |
| 167 | 693 | 3.77 | 3.42 | 0 | 2000 |
| 169 | 749 | 2.60 | 2.13 | 0 | 2000 |
| 172 | 787 | 3.51 | 3.08 | 0 | 2000 |
| 161 | 1063 | 5.45 | 1.52 | 1 | 0 |

**Figure S10.**
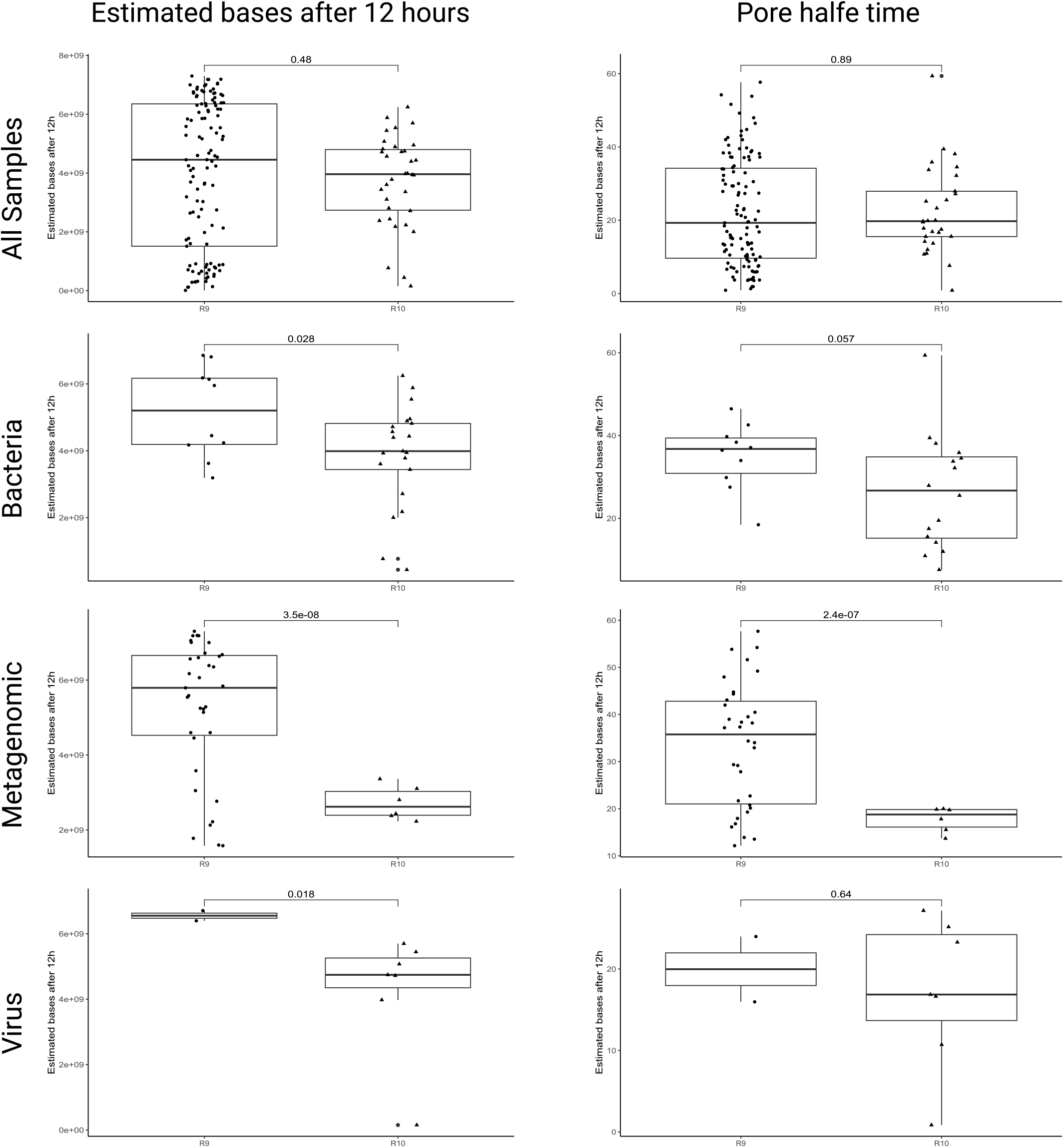
For the data shown in Fig. 6 B,C a t-test reveals no significant differences in estimated bases after 12 hours and pore half time between R9 and R10 data. However, when separating the data based on sample types, both EB12 and pore half time show significant differences for the metagenomic samples.

**Figure S11.**
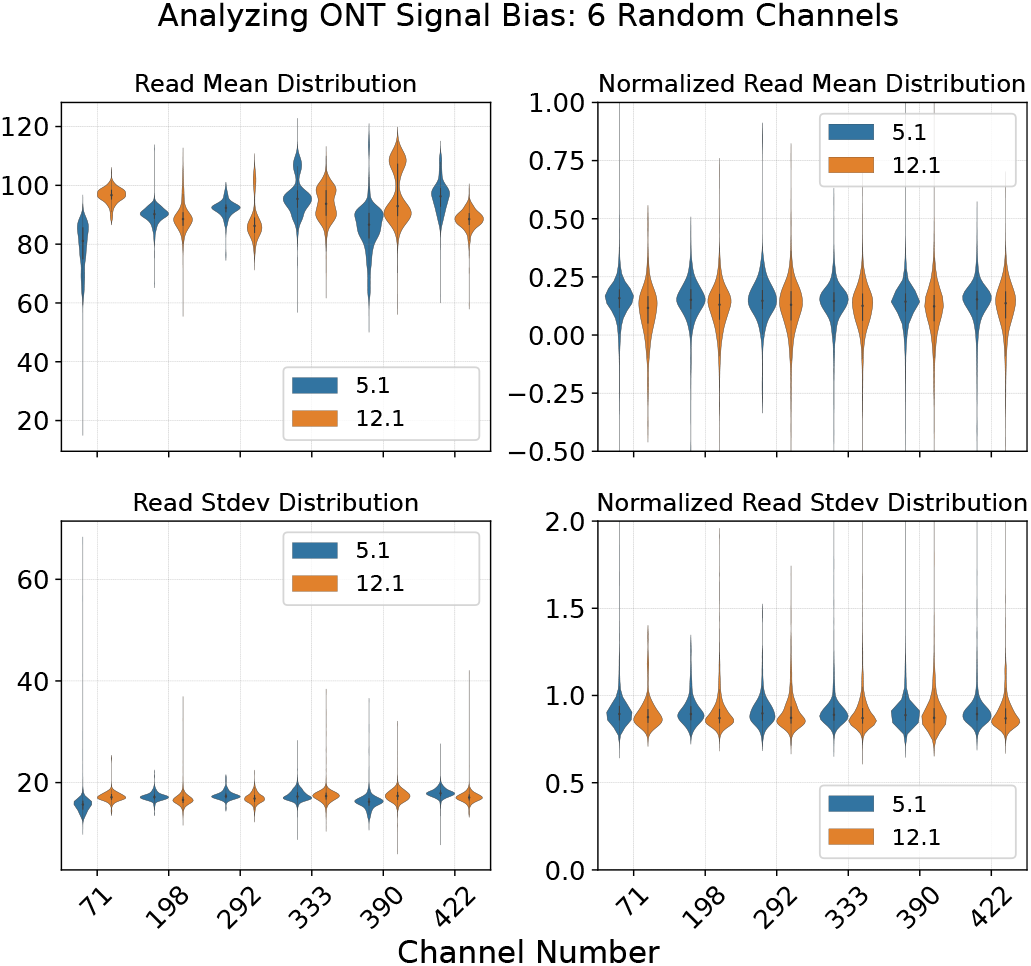
Read mean distribution (top) and read standard deviation (bottom) before (left) and after (right) normalization of six randomly picked channels from the sequencing runs 5.1 and 12.1. Biases caused by pores, flow cells, or sensors vanish after normalization.

## Footnotes

1 https://community.nanoporetech.com

2 https://nanoporetech.com/document/flow-cell-check

3 https://nanoporetech.com/document/hardware

4 https://nanoporetech.com/document/chemistry-technical-document

5 https://community.nanoporetech.com/attachments/10325/download

6 https://nanoporetech.com/document/requirements/lig-seq-input-load

7 according to ONT Live support asking “What is DCS?”, https://nanoporetech.com, 12.12.2024, see Fig. S1

8 according to ONT Live support asking “What is RCS?”, https://nanoporetech.com, 12.12.2024, see Fig. S2

9 Note: 1 *μ*g DNA of 8 kb fragments are 200 fmol; 5 kb are about 300 fmol; 50 kb are 30 fmol

10 https://international.neb.com/protocols/2012/11/01/protocol-for-use-with-end-user-supplied-primers-and-adaptors-e6000

11 https://nanoporetech.com/document/experiment-companion-minknow “Chapter 43. Troubleshooting your run from the pore activity plots”

12 https://github.com/artic-network/artic-ncov2019

13 23 October 2024

14 https://nanoporetech.com/document/adaptive-sampling

15 https://nanoporetech.com/document/adaptive-sampling

16 https://github.com/nanoporetech/dorado

17 https://github.com/nanoporetech/bonito

18 https://github.com/nanoporetech/remora

19 https://github.com/a-slide/pycoQC

20 https://github.com/wdecoster/NanoPlot

21 https://github.com/wdecoster/nanoQC

22 https://github.com/wdecoster/chopper

23 https://github.com/rrwick/Filtlong

24 https://github.com/lh3/minimap2

25 https://github.com/marbl/Winnowmap

26 https://github.com/paoloshasta/shasta

27 https://github.com/Nextomics/NextDenovo

28 https://github.com/nanoporetech/medaka

29 https://github.com/isovic/racon

30 https://github.com/rnajena/read5

31 https://github.com/nanoporetech/megalodon

32 https://github.com/nanoporetech/dorado

